# Profile of basal cell carcinoma mutations and copy number alterations - focus on gene-associated noncoding variants

**DOI:** 10.1101/2021.06.24.449728

**Authors:** Paulina Maria Nawrocka, Paulina Galka-Marciniak, Martyna Olga Urbanek-Trzeciak, M. Ilamathi, Natalia Szostak, Anna Philips, Laura Susok, Michael Sand, Piotr Kozlowski

## Abstract

Basal cell carcinoma (BCC) of the skin is the most common cancer in humans, characterized by the highest mutation rate among cancers, and is mostly driven by mutations in genes involved in the hedgehog pathway. To date, almost all BCC genetic studies have focused exclusively on protein-coding sequences; therefore, the impact of noncoding variants on the BCC genome is unrecognized. In this study, with the use of whole-exome sequencing of 27 tumor/normal pairs of BCC samples, we performed an analysis of somatic mutations in both protein-coding sequences and gene-associated noncoding regions, including 5’UTRs, 3’UTRs, and exon-adjacent intron sequences. Separately, in each region, we performed hotspot identification, mutation enrichment analysis, and cancer driver identification with OncodriveFML. Additionally, we performed a whole-genome copy number alteration analysis with GISTIC2. Of the >80,000 identified mutations, ∼50% were localized in noncoding regions. The results of the analysis generally corroborated the previous findings regarding genes mutated in coding sequences, including *PTCH1*, *TP53*, and *MYCN,* but more importantly showed that mutations were also clustered in specific noncoding regions, including hotspots. Some of the genes specifically mutated in noncoding regions were identified as highly potent cancer drivers, of which *BAD* had a mutation hotspot in the 3’UTR, *DHODH* had a mutation hotspot in the Kozak sequence in the 5’UTR, and *CHCHD2* frequently showed mutations in the 5’UTR. All of these genes are functionally implicated in cancer-related processes (e.g., apoptosis, mitochondrial metabolism, and *de novo* pyrimidine synthesis) or the pathogenesis of UV radiation-induced cancers. We also found that the identified *BAD* and *CHCHD2* mutations frequently occur in melanoma but not in other cancers via The Cancer Genome Atlas analysis. Finally, we identified frequent deletion of chr9q, encompassing *PTCH1*, and unreported frequent copy number gain of chr9p, encompassing the genes encoding the immune checkpoint ligands PD-L1 and PD-L2. In conclusion, this study is the first systematic analysis of coding and noncoding mutations in BCC and provides a strong basis for further analyses of the variants in BCC and cancer in general.

**Author summary:** The study is the first systematic analysis of both coding and noncoding mutations in basal cell carcinoma (BCC), the most common and the most highly mutated human cancer. Noncoding mutations accounted for ∼50% (∼40K mutations) of all mutations detected by the standard WES approach. Among the genes frequently mutated in noncoding regions are: *BAD* with a hotspot in the 3’UTR, *DHODH* with a hotspot in the Kozak sequence, and *CHCHD2* with mutations in 5’UTR, all genes functionally related to cell metabolism and apoptosis. Analysis of copy number alterations showed frequent chr9q deletion, encompassing *PTCH1* (the key BCC tumor suppressor), and frequent copy number gain of chr9p, encompassing the genes of the immune checkpoint proteins PD-L1 and PD-L2.

## INTRODUCTION

Basal cell carcinoma (BCC), a type of nonmelanoma skin cancer (NMSC), is the most common human cancer affecting predominantly elderly people of the Caucasian population [1–3]. The lifetime risk of BCC in the Caucasian population is ∼30%, and it is higher in men and fair-skinned people. BCC usually occurs sporadically but can also develop as a result of Gorlin syndrome (also known as nevoid basal cell carcinoma syndrome (NBCCS)), an autosomal dominant hereditary condition with an incidence of approximately 1:30,000 [4] characterized by the frequent appearance of multiple BCC lesions that develop at a younger age together with skeletal abnormalities, odontogenic keratocysts, and an increased risk of medulloblastoma. Histologically, BCCs are classified into three major subtypes: nodular, which is the most common subtype; superficial; and infiltrative or sclerodermiform. Other subtypes as well as mixed types occur less frequently [5–7]. Predominantly, superficial and nodular BCCs are slow-growing, locally invasive, epidermal tumors with a metastasis rate of <0.1% [8,9], while infiltrative BCCs are characterized by more aggressive, tong-like, subclinical growth patterns mimicking icebergs, as they often grow below clinically healthy-looking skin [10,11]. Although BCC aggressiveness overall is low and metastatic potential is negligible, the commonness of BCC and the increasing incidence associated predominantly with aging populations has brought attention to its pathogenesis [2,3,12–17]. Exposure to ultraviolet (UV) radiation, which can lead to point mutations frequently represented by C>T and CC>TT transitions, is the main causative factor in the pathogenesis of BCC [18]. Additional risk factors include ionizing radiation, arsenic ingestion, and immune suppression [19,20].

BCC is characterized by the highest mutation rate observed among cancers, having over 65 mutations/Mbp [14,15]. The most frequent genetic alterations occurring in BCC are mutations disturbing the hedgehog (SHH/PTCH1/SMO) pathway, predominantly loss-of-function mutations in *PTCH1* but also activating mutations in *SMO*; these genes encode two transmembrane proteins, PTCH1 (also known as Patched1) and SMO (also known as Smoothened), respectively [14,15]. The pathway is activated by the SHH signaling protein (also known as Sonic hedgehog), which binds to the extracellular domain of PTCH1, disabling inhibition of SMO; this in turn activates GLI transcription factors. Germline mutations in *PTCH1* predispose patients to Gorlin syndrome [21].

Previous studies, including whole-exome sequencing (WES) analyses, have also recognized other genes/pathways frequently mutated in BCC, including *TP53, MYCN, PPP6C, PTPN14, STK19,* and *LATS1* [14,15], as well as genes involved in the RTK-RAS-PI3K and Hippo-YAP pathways [15]. However, as an overwhelming majority of BCC genetic studies (as well as those in other cancers) have focused almost exclusively on protein-coding sequences, very little is known about mutations in noncoding regions (noncoding mutations). Noncoding mutations are not studied/reported even if detected, e.g., as a result of WES. On the other hand, it is well known that the noncoding parts of genes, i.e., promoters, introns, or 5’ and 3’ untranslated regions (5’UTRs and 3’UTRs, respectively), encompass numerous functional elements important for the proper functioning of the genes [22–24]. Somatic mutations may disrupt or modify the properties of these elements, acting either as gain- or loss-of-function mutations and thus enhancing/accelerating or switching off the function of some genes. Despite the negligence of the analysis of noncoding regions, there are some spectacular examples of noncoding driver mutations, for example, TERT promoter mutations, which occur most frequently in melanoma brain and bladder cancers but are also reported in BCC [25–27], and mutations in the precursor of miR-142, which frequently occurr in Hodgkin lymphomas and acute myeloid leukemia (summarized in [28]). The miRNA biogenesis enzyme DICER has also been shown to bear mutations that could play a role in aberrant miRNA expression in BCC [29–31]. It should also be noted that an effort to catalog cancer somatic mutations in the noncoding genome has recently been undertaken [32,33]; however, this pancancer project does not include BCC.

To preliminarily explore the occurrence of noncoding somatic mutations in BCC, we performed WES of over two dozen BCC samples, extending the analysis beyond protein-coding sequences and focusing on gene-associated noncoding regions, i.e., 5’UTRs, 3’UTRs, and exon-adjusted sequences of introns. Apart from the fact that our results well-replicate those of previous BCC studies in terms of mutations in protein-coding genes, we showed that a substantial portion of mutations are located in noncoding regions. Many of these mutations frequently recur in particular noncoding regions or in specific hotspot positions. Computational analyses showed that some of the gene mutations in noncoding regions are potential cancer drivers and are functionally related to skin cancers. Additionally, whole-genome copy number alteration (CNA) analysis revealed frequent deletion of chr9q, encompassing *PTCH1*, and unreported frequent amplification of chr9p, including the genes encoding two immune checkpoint ligands PD-L1 and PD-L2.

## RESULTS

### Overall sequencing and mutation occurrence characterization

We performed WES on 27 paired tumor and corresponding intraindividual control skin DNA samples isolated from 22 nodular and 5 superficial BCC subtypes and corresponding healthy skin tissue. The average coverage of the targeted regions was 183x (185x in normal and 180x in tumor samples), ranging in different samples from 134x to 232x. In total, we identified 84,571 cancer-sample-specific somatic mutations (S1 Table), of which 42,380 (50.1%) were located in protein-coding (coding) regions, and the remaining 42,191 (49,9%) were located in noncoding regions (Table 1, Fig 1A). The noncoding regions included (i) 5’UTRs, (ii) ∼100 bp fragments of 3’UTRs adjacent to coding sequences (3’UTRs), (iii) exon-adjacent ∼100 bp fragments of introns (introns), and (iv) sequences other than those classified above (i-iii), mostly intergenic sequences located upstream and downstream of the first and last gene exons (intergenic regions). The average coverage of the mutated positions was 169x and was slightly higher in coding (195x) than in noncoding regions (142x), whereas the average fraction of reads mapping to alternative alleles was 0.35 (0.33 in coding and 0.40 in noncoding regions). The average mutation rate in the targeted regions was 52.8 mutations/Mb (ranging from 0.1 to 287.5), which, although slightly lower than that observed before in BCC [15,34], is still higher than that in any other tested cancer type. Although somewhat counterintuitive, the lower mutation burden in our study than in other BCC studies [15,34] may result from the much higher sequencing coverage in our study, which gave us much higher statistical power to filter out the fraction of false-positive mutations. The lower mutation burden in our study may also be explained by the identification in our cohort of two samples with an extremely low mutational burden (<0.2 mutations/Mb). Most of the identified mutations were single-nucleotide substitutions (79,960 (94.5%), predominantly C>T transitions), followed by double substitutions (3,128 (3.7%), predominantly CC>TT transitions) and short (<4 nt) indels (1.483 (1.8%)) (Table 1, Fig 1B). The higher frequency of indels in noncoding regions most likely results from the excess of low complexity sequences, which cause polymerase slippage.

**Table 1.**
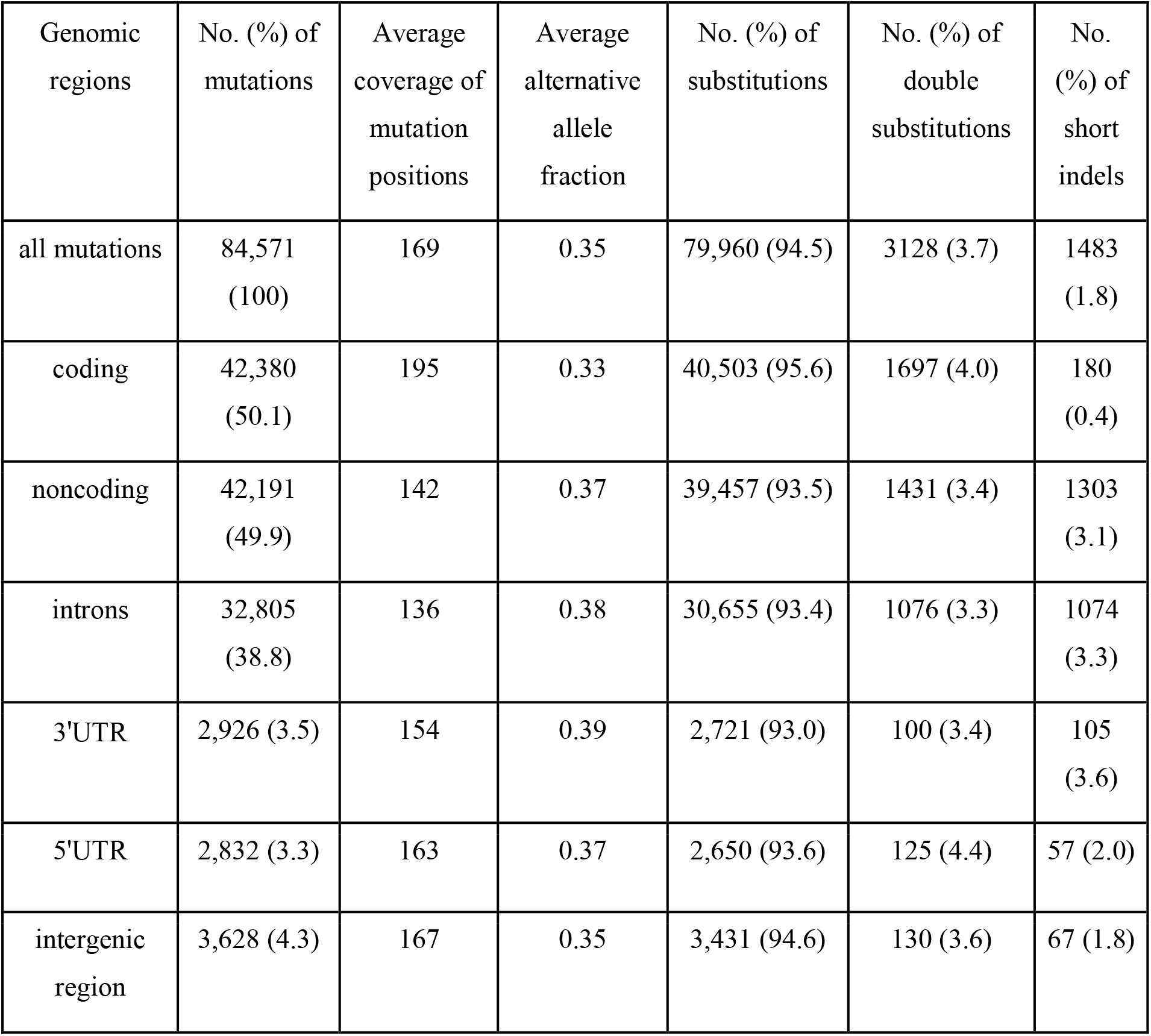
Summary of somatic mutation distribution and mutation types in BCC.

| Genomic regions | No. (%) of mutations | Average coverage of mutation positions | Average alternative allele fraction | No. (%) of substitutions | No. (%) of double substitutions | No. (%) of short indels |
| --- | --- | --- | --- | --- | --- | --- |
| all mutations | 84,571 (100) | 169 | 0.35 | 79,960 (94.5) | 3128 (3.7) | 1483 (1.8) |
| coding | 42,380 (50.1) | 195 | 0.33 | 40,503 (95.6) | 1697 (4.0) | 180 (0.4) |
| noncoding | 42,191 (49.9) | 142 | 0.37 | 39,457 (93.5) | 1431 (3.4) | 1303 (3.1) |
| introns | 32,805 (38.8) | 136 | 0.38 | 30,655 (93.4) | 1076 (3.3) | 1074 (3.3) |
| 3'UTR | 2,926 (3.5) | 154 | 0.39 | 2,721 (93.0) | 100 (3.4) | 105 (3.6) |
| 5'UTR | 2,832 (3.3) | 163 | 0.37 | 2,650 (93.6) | 125 (4.4) | 57 (2.0) |
| intergenic region | 3,628 (4.3) | 167 | 0.35 | 3,431 (94.6) | 130 (3.6) | 67 (1.8) |

**Fig 1.**
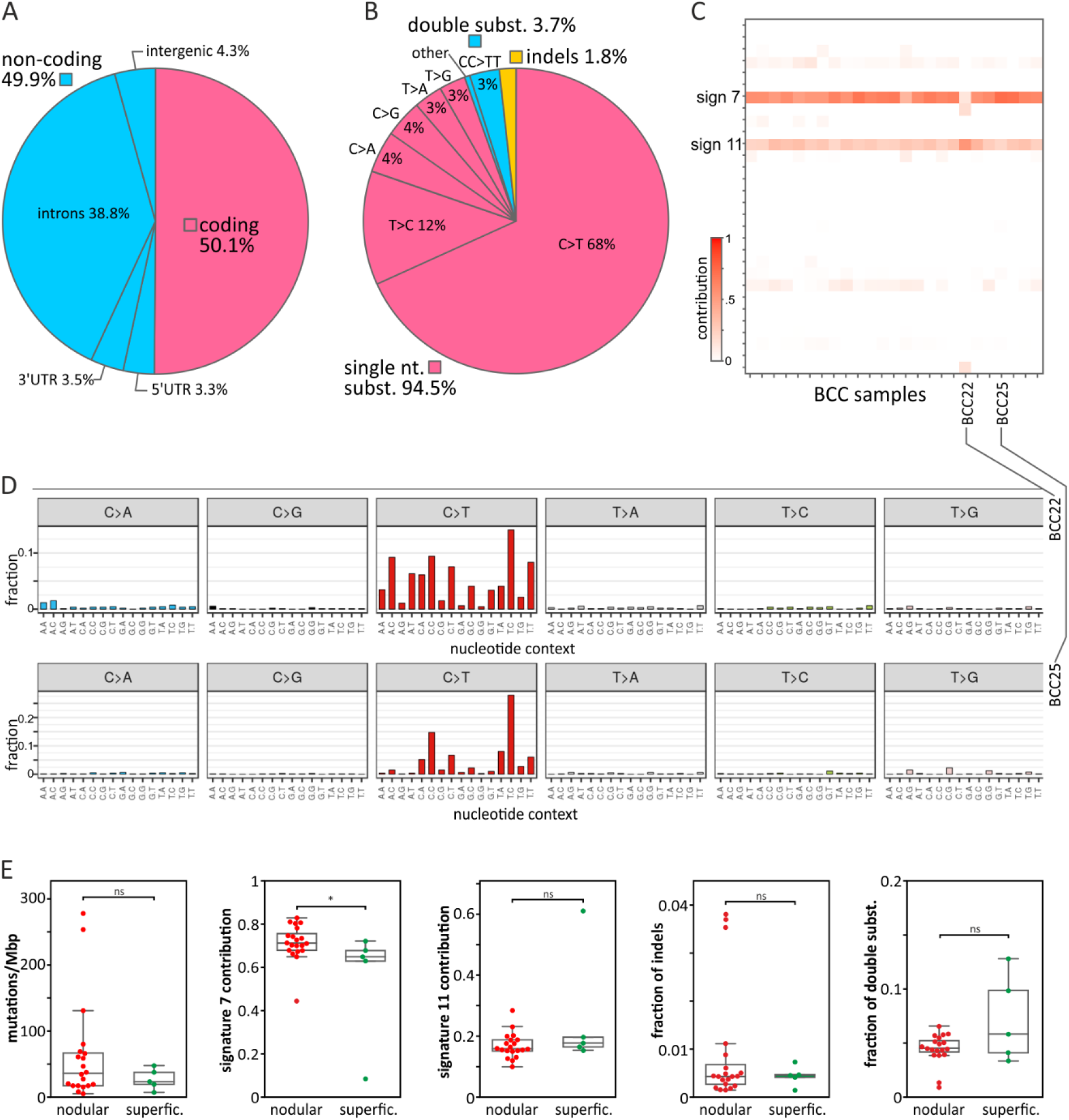
Mutation distribution, mutational signatures, and comparison of superficial and nodular BCC subtypes. A) Frequency of mutations in particular gene/genomic regions. B) Frequency of mutation types. C) Heatmap showing the contribution of the mutational signatures (rows) to the analyzed BCC samples (columns). A higher color intensity indicates a higher contribution (as indicated on the scale bar). D) Representative mutation distribution plots of samples with a high association with signature 7 (sample BCC25) and signature 11 (sample BCC22). E) Comparison of nodular and superficial BCC samples in terms of (from the left) mutational load, signature 7 and signature 11 contributions, frequency of indels, and frequency of double substitutions.

To estimate the fraction of false-positive mutations, we resequenced (with Sanger sequencing) 52 mutations representing different types of alterations, including 39 substitutions and 13 indels (S2 Table). The analysis confirmed 51/52 of the mutations, indicating a very low (2%) fraction of false-positive mutations. The fraction may be even lower, as the only unconfirmed mutation (double substitution CC>TT in *MYCN*) was present in a low fraction of reads (7%), which is generally beyond the sensitivity of Sanger sequencing.

### Mutational signatures

The analysis of mutational signatures showed that most of the samples were predominantly associated with signature 7 (average signature contribution (SC) = 0.7) and to a lesser extent with signature 11 (average SC = 0.2) (Fig 1C-D). Both signatures consist predominantly of C>T substitutions but differ in the sequence context of the substitutions. Signature 7 is associated with UV irradiation exposure and commonly occurs in melanoma and head and neck cancer. A hallmark of signature 7 is the frequent occurrence of double CC>TT substitutions resulting from UV radiation-induced pyrimidine dimers. Signature 11 was previously found in melanoma and glioblastoma multiforme, often in patients treated with the alkylating agent *temozolomide*, which is also used in BCC therapy. Only one sample (BCC22) showed a stronger association with signature 11 (SC = 0.6) than signature 7 (SC = 0.3). None of the analyzed samples showed an association with signatures 1, 2, 5, and 13, which are frequent in most cancer types. This may indicate that the deamination of 5-methylcytosine (5meC) predominantly induced by AID/APOBEC cytidine deaminases (attributed to the abovementioned signatures) does not play a role in the pathogenesis of BCC.

The comparison of the nodular and superficial BCC samples showed no substantial difference in terms of mutation burden or mutation types, with the exception of the contribution to mutational signature 7, which was higher for the nodular than superficial samples (Fig 1E), consistent with the higher UV radiation exposure of nodular BCCs.

### Hotspot mutations

As recurrent mutations may be indicators of the cancer-related function of the mutated genes, we first looked for hotspots defined as genomic positions mutated in at least 3 samples (>10% of the cohort). In total, we identified 43 hotspots, including 23 hotspots in coding and 20 hotspots in noncoding regions (8 in 5’UTRs, 1 in 3’UTRs, and 11 in introns) (S3 Table). Of the coding hotspots, 16 resulted in missense mutations, and 7 were synonymous substitutions. Although the functionality of individual synonymous mutations cannot be unequivocally ruled out [35–37], the majority of these hotspot mutations resulted from randomly occurring neutral alterations, so we did not analyze them further. As shown in S3 Table, some of the hotspots were located in genes annotated in the COSMIC Cancer Gene Census (CGC) database and/or in genes playing a role in cancer or skin function.

#### Hotspot mutations in coding regions

Of the coding mutations (S3 Table), the most commonly identified in our study (in 5 samples) was the c.1292C>T (Ser431Phe) substitution, located at chr14:103,131,144 in the Sec6 domain of *TNFAIP2*, which encodes a multifunctional protein playing a role in angiogenesis, inflammation, cell migration and invasion, cytoskeleton remodeling, and cell membrane protrusion formation [38–41]. Nonetheless, *TNFAIP2* is not well-recognized in cancer, and the hotspot or other mutations in the gene have not been reported before. Another coding hotspot, mutated in 3 samples with the c.655C>T (Pro219Ser) substitution, was located at chr7:148,827,237 in *EZH2*; *EZH2* encodes an essential subunit (methyltransferase) of polycomb repressive complex 2 (PRC2), which plays a role in histone methylation and gene silencing [42]*. EZH2* is a well-known oncogene associated with a more aggressive form and poorer prognosis of many cancers, including melanoma, squamous cell carcinoma (SCC), and BCC, with demonstrated increased expression in SCC (compared to normal skin and SCC precursor actinic keratosis (AK)) [43] and aggressive BCC [44]. Both gain- and loss-of-function mutations in *EZH2* have often been found in myeloid leukemias and lymphomas but are not common in solid tumors. Contrary to the previously detected mutations clustering mostly in the catalytic SET domain (Bödör et al. 2013; Donaldson-Collier et al. 2019), the hotspot detected here was located in the N-terminal (NT) part of the protein, which, among other areas, is responsible for interaction with histones [47]. Whether the mutations may affect the interaction warrants further investigation. To the best of our knowledge, this mutation hotspot has not been observed in any cancer, including BCC.

An additional interesting coding hotspot (mutated in 3 samples) was located at chr15:40,382,906-40,382,907. The hotspot was mutated with either the c.71C>T substitution or the c.71_72delinsTT double substitution (note that double substitutions are annotated as deletion/insertion (delins) variants according to HGVS nomenclature), both resulting in the Ser24Phe missense mutation affecting the NT part of the KNSTRN protein (also known as small kinetochore-associated protein (SKAP)), which plays a role in maintaining chromatid cohesion and proper chromatid separation during anaphase [48]. *KNSTRN* mutations (predominately the Ser24Phe hotspot mutation) were first detected in 19% of SCCs and 13% of AKs [49]. Subsequent analysis of The Cancer Genome Atlas (TCGA) datasets showed that the *KNSTRN* mutations also occur in 5% of melanoma samples but are rare in other cancers. Later, *KNSTRN* mutations were also identified in 2% [15] and 10% [50] of BCCs. These findings together with this study confirm that *KNSTRN* mutations are specific to UV radiation-related skin cancers. Consistent with the role of KNSTRN, it was shown that *KNSTRN* mutations in SCC affect proper chromosome separation and are associated with increased chromosome instability, expressed as a fraction of the genome with copy number alterations (CNAs) [49]. Although there was a similar number of tested samples, the association of the KNSTRN mutations with CNAs was not confirmed in BCC, neither in a study by Jaju et al. [50] nor in our study (S1 Fig). It is worth noting that it was also shown that KNSTRN plays a role in UV radiation-induced apoptosis [51]; however, the effect of the mutations on avoidance of apoptosis by BCC cells or any other cancer cells has not yet been tested.

#### Hotspot mutations in noncoding regions

The most frequently mutated hotspot of all the hotspots detected in the study hotspot (mutated in 8 samples) was located at chr11:64,270,066-64,270,067 in the 3’UTR of *BAD* and has never been reported before. The hotspot encompasses 4 different substitutions (c.*142C>A, c.*142C>T, c.*142_*143delinsTT and c.*143C>T; S3 Table and Fig 2A), located 142 or 143 nucleotides (nt) downstream of the stop codon. The protein encoded by the gene is a member of the BCL-2 family, which plays a role in the positive regulation of cell apoptosis. The gene is commonly implicated in many cancers [52,53]; however, to the best of our knowledge, this hotspot has not been reported before in any cancer.

**Fig 2.**
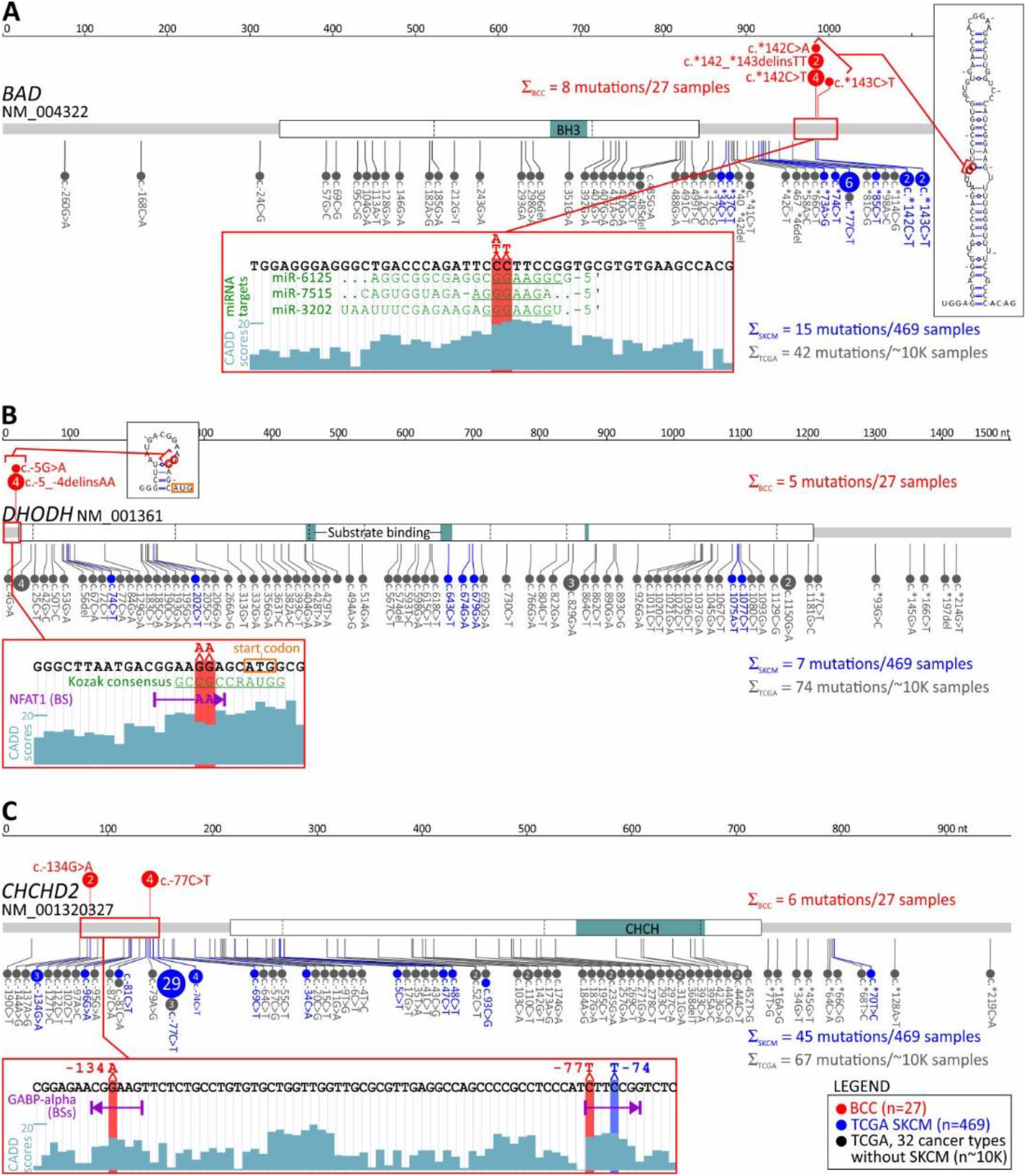
Distribution of mutations in the selected genes with the identified mutation hotspots in noncoding areas. A)-C) Maps of the *BAD*, *DHODH*, and *CHCHD2* genes, with the exon structure and protein functional domains indicated. Mutations are visualized in the form of lollipop plots along with the gene maps, and the size of a mutation symbol (circle) is proportional to the number of mutations. Mutations identified in BCC (red) are shown above and mutations identified in SKCM (blue) and other TCGA cancers (gray) are shown below the maps. The inset below each map shows the detailed sequence context of the hotspot mutations, along with CADD score graphs, indicating the functional relevance of particular positions and other sequence characteristics (i.e., (in A) predicted miRNA target sites, (in B) the Kozak consensus sequence and NFAT1 transcription factor binding sites (BSs) created by the hotspot mutation, and (in C) the GABP-alpha transcription factor BSs disrupted by the hotspot mutations). The additional insets in A and B show computationally predicted RNA secondary structures generated from RNA sequences directly flanking the hotspots.

Next, another novel noncoding hotspot mutated in 5 samples located at chr16:72,008,760-72,008,761 in the 5’UTR of *DHODH* was identified. The hotspot encompasses two different substitutions, c.-5G>A and c.-5_-4delinsAA, affecting the Kozak sequence (S3 Table and Fig 2B). *DHODH* is not well studied in cancer, but it has recently been demonstrated that it plays an important role in the carcinogenesis of SCC and other UV radiation-induced skin cancers [54,55].

Another mutated noncoding hotspot from our study worth mentioning was found in 4 samples with the c.-77C>T substitution and was located at chr7:56,106,490 in the 5’UTR of *CHCHD2*, also known as *MNRR1* (S3 Table and Fig 2C). The analysis of the entire *CHCHD2* 5’UTR showed one more recurrent (in 2 samples) substitution, c.-134G>A, located at chr7:56,106,547, resulting in a total of 6 mutations in the 5’UTR in 6 samples. Interestingly, frequent mutations in the hotspot in the 5’UTR of *CHCHD2* were previously reported in melanoma [56].

Finally, we identified a hotspot located at chr1:153,990,763 in the 5’UTR of *RPS27* (encoding a ribosomal protein component of the 40S subunit) that was mutated in 3 samples with the c.- 34C>T substitution. Mutations in the promoter/5’UTR of *RPS27* (including the hotspot mutation) have been identified before in ∼10% of melanoma samples [56,57] but have never been reported in BCC or other skin cancers. Subsequent *in vitro* functional studies showed that the *RPS27* 5’UTR hotspot mutation decreases *RPS27* mRNA levels and that decreased levels of RPS27 are associated with worse prognosis of melanoma patients and drug (*vemurafenib* and *palbociclib*) sensitivity of melanoma cells [58].

#### Computational analysis of the identified noncoding hotspots and comparison with external datasets

To further characterize three noncoding hotspot mutations, two not previously reported in *BAD* and *DHODH* and one in *CHCHD2* previously reported in melanoma [56], we analyzed their potential impact with a number of computational tools and investigated their incidence in other cancers using external datasets of a large cohort (>10,000 samples) of TCGA samples, representing 33 different human cancer types (including 469 skin cutaneous melanoma (SKCM) samples but not including BCC or SCC samples). Note that the list and the standard abbreviations of all TCGA cancer types are in S4 Table.

In total, in the TCGA samples, we identified 28 mutations in the *BAD* 3’UTR (Fig 2A). The mutations were found predominantly in SKCM samples(15 mutations in 12 (2.6%) SKCM samples), including 4 mutations in the hotspot (residues c.*142C and c.*143C) identified in BCC, and 6 c.*77C>T mutations, constituting an additional hotspot in the 3’UTR, not occurring in BCC. In other cancers, 3’UTR mutations were very rare (Fig 2A). In contrast with the mutation frequency in the 3’UTR, mutations in other parts of the gene, including the coding region (n=26, predominantly missense or synonymous), were rare (not exceeding 1% in any cancer) and randomly distributed between different cancer types (excluding SKCM). The exclusiveness of the SKCM and BCC mutations in the 3’UTR vs. other parts of the gene (enrichment compared to other cancer types; Fisher’s exact test; p<0.0001 and p=0.0005, respectively) precludes an accidental occurrence of the mutations, solely as a result of some region- and/or mutagenesis-related mechanisms and argues for the cancer-driven selection of the 3’UTR mutations in BCC and SKCM (and likely also in other UV irradiation-related cancers).

Next, with the use of TargetScan, we identified 3 miRNAs (miR-7515, miR-3202, and miR-6125) whose predicted targets (seed-interacting sequences) were disrupted by hotspot mutations (Fig 2A). However, as (i) none of these targets has been validated by any means (miRTarBase; [59]), (ii) none of these miRNAs have been confidently validated (via miRBase or miRGeneDB), and (iii) none of these miRNAs have been found to have expression levels detectable/confirmed in any of the TCGA cancers, it is very unlikely that any of the identified targets are functional. Additionally, the occurrence of SKCM mutations in different positions across the *BAD* 3’UTR argues against the possibility that the driving force of the mutations is a disruption of a particular miRNA target. Some clue for the functionality of the BCC hotspot may be its location in the 5’ arm of the ∼40 bp long stable hairpin RNA structure motif (dG=-39.6 Kcal/mol), which is destabilized (by ∼2 Kcal/mol) by the hotspot mutations (Fig 2A).

The analysis of TCGA data showed no mutation in the BCC hotspot or any other mutation in the *DHODH* 5’UTR in any of the TCGA cancer types, even though different mutations (n=81) were identified in other parts of the gene, including 75 mutations in the coding region (Fig 2B). The other mutations, however, were randomly distributed along the gene sequence and between different cancer types, and only two of the coding mutations were deleterious (frameshift) mutations. This result indicates that the *DHODH* 5’UTR hotspot mutations are BCC-specific mutations, and the absence of these mutations in other UV radiation-related cancers makes it unlikely that the frequent occurrence of the mutations in BCC is solely due to a random effect of UV irradiation. The 5’UTR of *DHODH* is very short (21 bp). Although hotspot mutations occurred in the Kozak sequence, which is important for the initiation of translation, neither wild-type nor mutant alleles affected the consensus Kozak sequence nucleotides (at positions - 4 and -5); therefore, the ATGpr [60]), and NetStart 1.0 (GedersenAG [61] tools predicted the mutations to have a minor effect on the effectiveness of translation under standard conditions. However, this result does not exclude an effect of the mutations under specific conditions, such as hypoxia, UV exposure, or cancer.

The analysis of RNA secondary structure showed that the hotspot mutations slightly modified (decreased the stability of) a small hairpin motif predicted to be formed by an RNA sequence directly flanking the hotspot (Fig 2B). The mutation may also destabilize the potential long-range interaction of the sequence flanking the mutations with the sequence located ∼200 nt downstream. Analysis of the 5’UTR sequence [62] showed that the double substitution (GG>AA) at the hotspot creates a consensus binding site for the NFAT1 transcription factor (Fig 2B), which is expressed in many tissues, including sun-exposed and non-sun-exposed skin (GTExPortal; GTEx Consortium Science 2020), and implicated in many cancers, including melanoma [63,64].

In total, in TCGA data, we identified 63 mutations in the *CHCHD2* 5’UTR (Fig 2C). The mutations were found predominantly in SKCM samples (40 mutations in 39 (8.5%) samples), including 29 c.-77C>T mutations and 3 c.-134G>A mutations, located in the hotspot positions identified in BCC. Additionally, we identified 4 samples with the c.-74C>T mutation, constituting an additional hotspot in the 5’UTR. Only 5 SKCM mutations were located outside the 5’UTR, 4 in the CDS (2 missense and 2 synonymous) and 1 in the 3’UTR (one mutation) (Fig 2C). In other cancers, there were rare 5’UTR mutations, including 4 mutations in HNSC and UCEC, 3 mutations in BRCA, and 12 mutations in other cancers. Three of these mutations coincided with the c.-77 hotspot. The positions of BCC/SKCM hotspot mutations seem to be nonrandom because they were all located in and all disrupted two distinct GABP-alpha transcription factor binding sites (mapped with the use of MotifMap [65]) (Fig 2C).

### Frequently mutated genes

Next, we looked at the overall frequency of mutations in the genes, separately analyzing mutations in coding regions, 5’UTRs, 3’UTRs, and introns (defined in Materials and Methods; listed in S5 Table). Although they were not considered frequently mutated, in this section, we also report genes with any mutations in a coding region if they were detected in a pathway of a recurrently mutated gene. In the analysis of frequently mutated regions, we focused mostly on genes functionally related to cancer (annotated with CGC and a manual literature search) and genes playing a role in skin function.

#### Genes frequently mutated in coding regions

In total, we identified 606 genes frequently mutated in coding regions. The most frequently mutated was *PTCH1*, with a total of 24 mutations in 20 BCC samples, including 5 missense, 4 splice-site, and 15 deleterious (nonsense or frameshift) mutations (Fig 3A). Mutation c.3450-1G>A located upstream of exon 21 was one of the splice-site mutations and was also observed in another study [14], which suggests its recurrence in BCC. We tested and confirmed the exon-skipping effect of the mutation with the use of exon-junction PCR and Sanger sequencing analysis (Fig 3A). The other genes from the hedgehog pathway recurrently mutated in our cohort were *GLI2*, which was mutated in 5 samples, and *SMO*, which was mutated in 4 samples (S4 Fig and Fig 4). The combined frequency of *SMO* and *GLI2* mutations was much lower in samples with (4/20; 20%) than in those without (4/7; 57%) *PTCH1* mutations, which suggests mutual exclusiveness of these mutations (Fig 4). Altogether, 24 (88%) samples had mutations in genes involved in the hedgehog pathway. Other frequently mutated cancer-related genes were *TP53* (7 missense, 8 deleterious, and one splice-site mutation in 13 samples) (Fig 3B); *MYCN* (8 missense mutations in 8 samples), *NOTCH1* (8 missense and 2 deleterious mutations in 8 samples), *NOTCH2* (3 missense, 3 deleterious, and 2 splice-site mutations in 7 samples), *NOTCH3* (6 missense mutations in 5 samples; note that the *NOTCH* mutations colocalized with the regions of the loss-of-function mutations identified in other solid tumors, e.g., in SCCs [66], *LATS1* (5 missense and one deleterious mutation in 5 samples), and *ARID1A* (5 missense mutations in 5 samples) (Fig 4 and S2 Fig). The mutations in the abovementioned genes are generally consistent with mutations observed before in BCC [14,15]. Additionally, we identified very frequent mutations (18 missense and 1 deleterious) in *PTPRD* (Fig 3C), a tumor suppressor frequently mutated in many cancers, including melanoma and cutaneous SCC [67–71], in 13 samples, but these have never been reported as frequently mutated in BCC. Interestingly, in addition to mutations in *MYCN*, we also noticed recurrent (although not frequent) mutations in three other genes in the MYC/MTOR regulatory network, i.e., *MTOR*, *DYRK3*, and *AMBRA1* (Fig 4), which have not been reported as mutated in BCC. The MTOR missense/activating mutations identified in other cancers are considered biomarkers for therapy with mTOR pathway inhibitors [72]).

**Fig 3.**
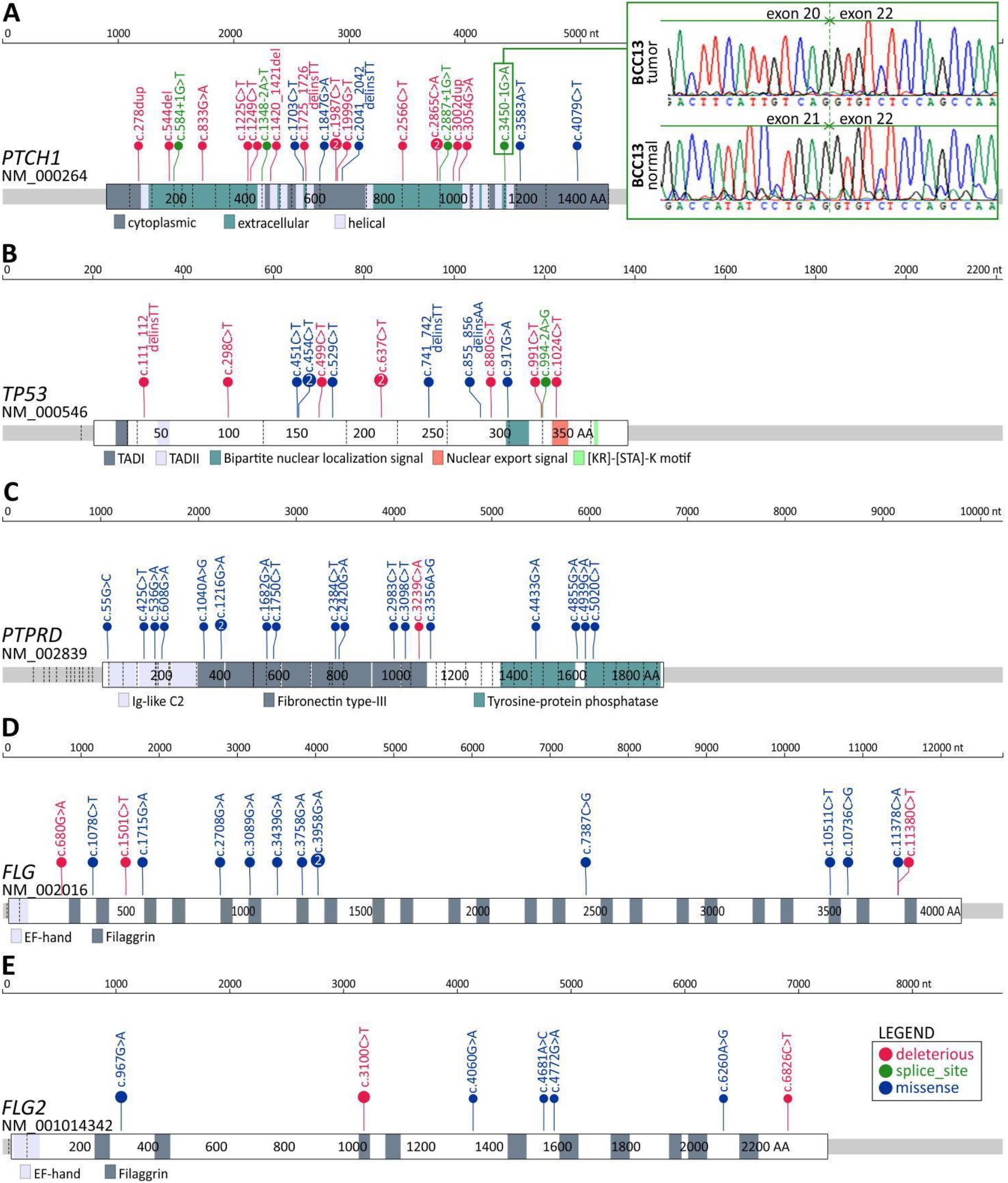
Distribution of the identified mutations in the genes with frequent mutations in the coding sequence. A)-E) Maps of the *PTCH1*, *TP53*, *PTPRD*, *FLG*, and *FLG2* genes. Mutations are visualized in the form of lollipop plots along with gene maps; the size of a mutation symbol (circle) is proportional to the number of mutations, and the color indicates the type of mutation (as shown in the legend). Additionally, the inset in A) shows the Sanger sequencing reads depicting the effect of the splice-site mutation c.3450-1G>A on exon 21 skipping.

**Fig 4.**
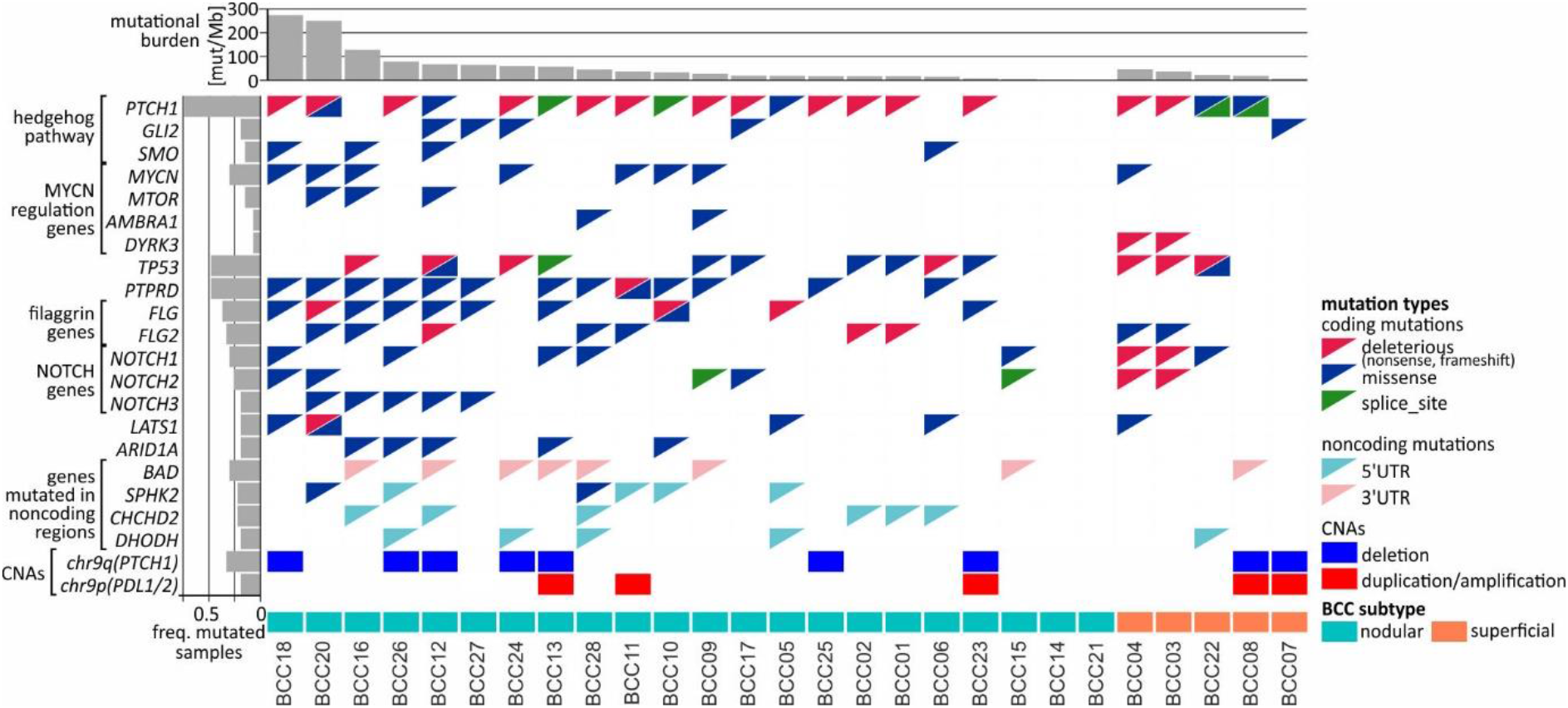
Comutation plot summarizing the somatic alterations in the BCC samples. Columns correspond to the samples, and rows correspond to the selected genes. The color of the mutation presence symbols corresponds to the mutation type, as indicated in the legend on the right. The bar plots above and on the left indicate the mutational burden and the fraction of samples with mutations in particular genes, respectively. The nodular and superficial samples are indicated by color.

Finally, we found a high frequency of mutations in the *FLG* (15 mutations in 10 samples) and *FLG2* (9 mutations in 9 samples) genes (Fig 3D-E and Fig 4), encoding profilaggrin and filaggrin-like proteins, precursors of filaggrin. Filaggrin is an important component of the stratum corneum of the epidermis that plays a role in maintaining epithelial homeostasis and barrier functions [73] and is a substrate for trans-urocanic acid (UCA) and pyrrolidone carboxylic acid (PCA), which are suggested to serve as a natural UV radiation barrier [74]. Although frequent mutations in the *FLG*/*FLG2* genes have been previously observed in other cancers, the mutations were usually considered random (passenger). Here, however, we observed a relatively high proportion of deleterious nonsense mutations, altogether occurring in 6 samples. Additionally, the analysis of the entire cohort of TCGA samples showed that the frequency of the *FLG*/*FLG2* mutations observed in our study in BCC substantially exceeds the frequencies of the mutations in other cancers, including melanoma (the next most frequently mutated cancer) (S3 Fig).

#### Genes frequently mutated in noncoding regions

Among the 11 genes frequently mutated in the 5’UTR (S5 Table) there were *DHODH* and *CHCHD2* with the hotspot mutations, as described in the previous paragraph. Of interest may also be *SPHK2*, with 4 dispersed mutations in 4 samples, whose function as both a proapoptotic gene suppressing cell growth and an oncogene promoting cell proliferation has been proposed [75–79]. *SPHK2* also had mutations in its coding region (Fig 4).

Among the 11 genes frequently mutated in the 3’UTR (S5 Table), in addition to *BAD*, which was described in the previous paragraph, we also identified 8 mutations in the 3’UTR of *SMIM27* (also annotated as lncRNA *TOPORS-AS1*); the overexpression of *SMIM27* was found to be associated with favorable outcomes in breast cancer [80].

Finally, we identified 289 genes (15 annotated in CGC) frequently mutated in introns (S5 Table). Interestingly, among the genes was *PTCH1*, which, in addition to 4 splice-site mutations (mentioned above), also had other 4 intronic mutations (in total, 8 intronic mutations). Other genes with frequent mutations in introns included *PTPRD* (14 mutations in 9 samples), which also frequently had mutations in the coding region; *NOTCH2* (6 mutations, including 2 splice-site mutations in 6 samples), which also frequently had mutations in the coding region; *ERBB4* (6 mutations in 6 samples), a well-known oncogene playing a role in many cancers (reviewed in [81]); and *DROSHA* (5 mutations in 5 samples), which encodes a core enzyme (nuclease) of the miRNA processing pathway and has been shown to be upregulated in BCC [82].

### Driver genes in BCC (OncodriveFML analysis)

To further investigate the mutations/mutated genes, we used OncodriveFML, which allows the prediction of the cancer driver potential of both coding and noncoding regions/genes based on functional mutation (FM) bias [83]. As shown in Fig 5A-C and S6 Table, we identified 14 potential cancer driver genes based on mutations in coding regions (CDS-drivers), a disproportionately high number of 36 potential cancer driver genes based on mutations in 5’UTRs (5’UTR-drivers), and 7 potential cancer driver genes based on mutations in 3’UTRs (3’UTR-drivers). No potential cancer driver gene was identified based on the mutations in introns.

**Fig 5.**
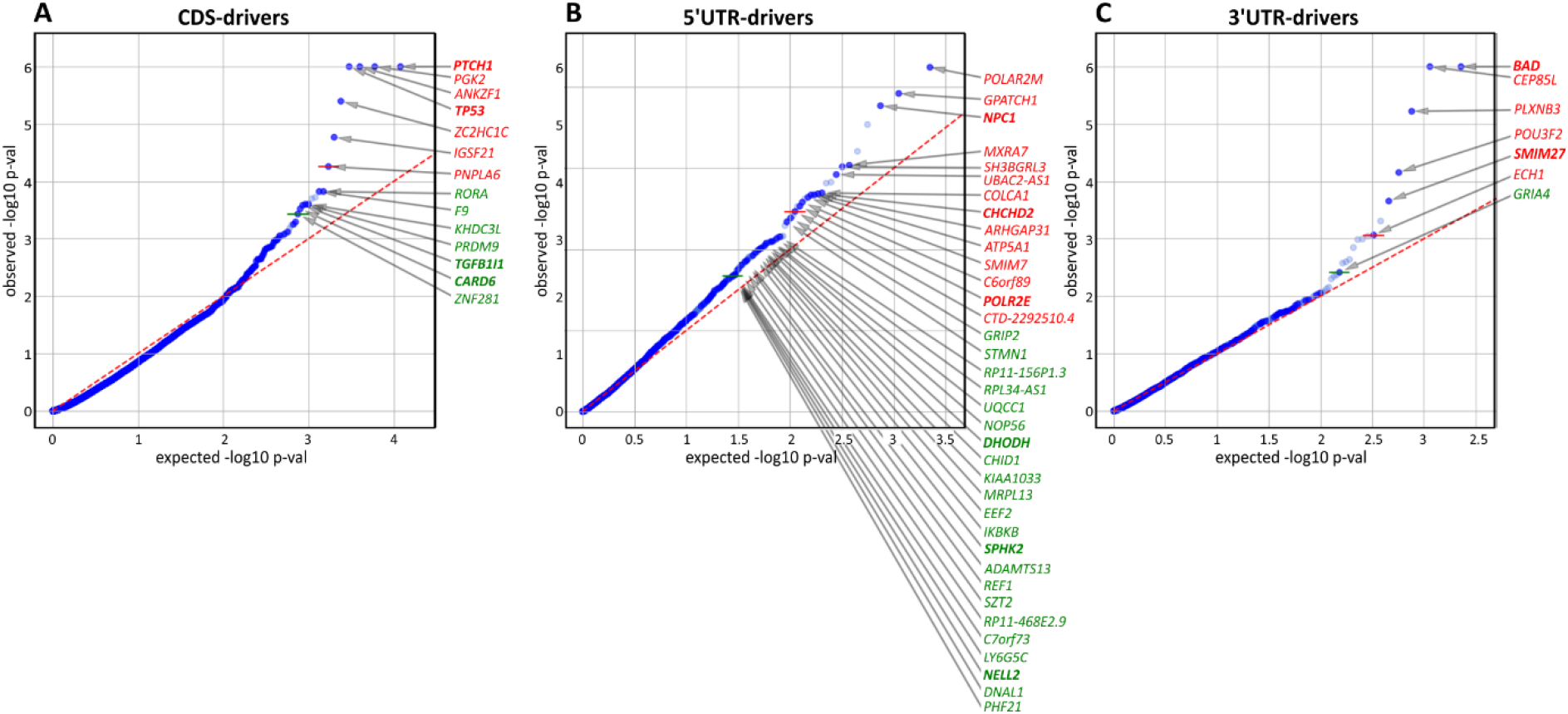
Identification of potential cancer drivers with the use of OncodriveFML. The quantile-quantile (QQ) plots show the distribution of expected (x-axis) and observed (y-axis) p-values corresponding to FM bias calculated (with CADD score) separately for mutations in A) coding regions, B) 5’UTRs, and C) 3’UTRs. The green and red colors indicate genes defined as significant (q<0.025) and highly significant (q<0.01), respectively, according to OncodriveFML recommendation.

In addition to 4 CDS-drivers (*PTCH1*, *TP53*, *TGFB1I1*, and *CARD6*) also identified as frequently mutated, it is worth noting *RORA*, recently shown to play an important role in restraining allergic skin inflammation [84]. Other interesting genes were *PRDM9* and *ZNF281*, both of which play a role in DNA repair and have been shown to be responsible for frequent mutations in cancer [85,86]. None of these genes were previously implicated or identified as frequently mutated in BCC.

Among the 5’UTR-drivers, 6 were also identified as frequently mutated: *DHODH*, *CHCHD2*, and *SPHK2* (described above), as well as *POLR2M*, *NPC1,* and *NELL2*. Additionally, it is worth noting *IKBKB* (mutated in 3 samples but not reported before as mutated in BCC) shown to act as a tumor suppressor in nonmelanoma skin cancers and noncancerous skin lesions; it was also shown that deletions of the gene lead to skin inflammation, hair follicle disruption, hyperplasia, and SCC development [87–90].

Among 3’UTR-drivers, two genes (mentioned above), i.e., *BAD* (the most significant 3’UTR-driver) and *SMIM27* were also identified as frequently mutated. Additionally, it is worth mentioning the transcription factor gene *POU3F2* (mutated in 3 samples), that plays a role in the invasiveness and metastasis of melanoma, and is controlled by miR-211 [91,92] and miR-107 [93]. Although the mutations were not located in the predicted miR-107 and miR-211 binding sites, they may affect the structure of the 3’UTR and thus indirectly change accessibility to these or other miRNA targets.

### Analysis of copy number alterations

As somatic CNAs have not been extensively studied in BCC, in the next step, we performed analysis of both chromosome arm-level and focal CNAs (with GISTIC2; [94]). At the chromosome arm level, we detected a significant recurring deletion of chr9q (q=1.4×10-6; occurring in 9 samples), involving *PTCH1* (Fig 4 and Fig 6), and a significant recurring amplification of chr9p (q=0.05; occurring in 5 samples), involving a region with *CD274* (also known as *PDL1*, encoding PD-L1), *CD273* (also known as *PDL2*, encoding PD-L2), and *JAK2* (Fig 4 and Fig 6).). Although the loss of chr9q has been frequently observed in BCC (reported as loss-of-heterozygosity of *PTCH1*), gain of chr9p has been reported only in one case of rare metastatic BCC [95]. To validate the chromosome 9 CNAs, we developed a multiplex ligation-dependent probe amplification (MLPA) assay with probes covering the entire chromosome 9 but especially focusing on the region containing *PTCH1* (chr9q22.32) and the region harboring *PDL1*, *PDL2*, and *JAK2* (chr9p24.1) (Fig 6). The MLPA analysis confirmed CNAs in all tested samples as detected by GISTIC2, and examples are shown in Fig 6.

**Fig 6.**
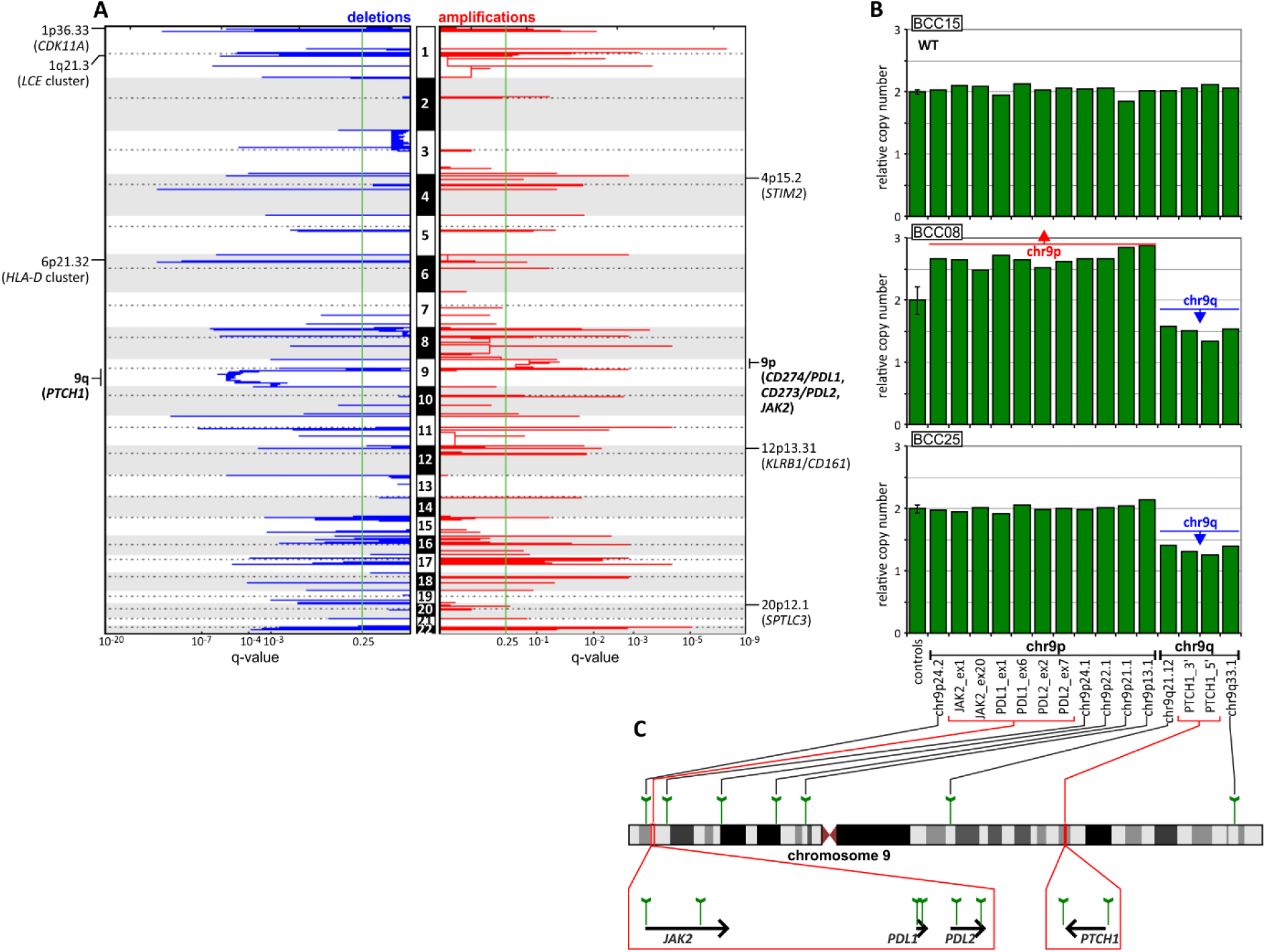
CNA analysis of the BCC samples. p A) GISTIC-estimated q-values for deletions (left, blue) and amplifications (right, red) are plotted along with chromosome positions (vertically). The green line indicates the recommended significance threshold, q=0.25. The selected significantly deleted and amplified regions/genes are indicated on the graphs. B) Representative MLPA results (bar plots), showing samples with chromosome 9 CNAs, i.e., chr9q deletion and chr9p amplification, vs. a sample (at the top) with the wild-type (WT) copy number genotype. Each bar plot depicts relative copy number values (y-axis) of the probes specific for regions along chromosome 9 and an average (with standard deviation error bar) signal of control probes (x-axis). C) Schematic depictions of the localization of the probes on chromosome 9 and in genes of interest.

CNA analysis also showed 54 regions of significant focal deletions, including 27 regions containing skin/cancer-related genes, and 56 significant amplifications, including 20 encompassing skin/cancer-related genes (Fig 6 and S7 Table). The elements involved in the most significant focal deletions were *CDK11A* (chr1p36.33; q=2.4×10-5; occurring in 6 samples), whose loss induces skin carcinogenesis [96]; the *LCE* cluster (chr1q21.3; q=2.4×10-6; occurring in 4 samples), including genes such as *LCE2* and *LCE3*, which play a role in maintaining skin barrier function and whose deletion has been associated with psoriasis [97]; and the *HLA-D* cluster (*HLA-DP*, -*DQ*, and *-DR*, chr6p21.32; q=2×10-4; occurring in 3 samples), encoding components of major histocompatibility complex (MHC) class II molecules, whose increased expression has been associated with increased cancer immunogenicity and better prognosis in BCC, SCC and melanoma [98–104]. The skin/cancer-related genes in the most significant focally amplified regions worth mentioning are *STIM2* (chr4p15.2; q=0.16; occurring in 2 samples) [105], *KLRB1/CD161* (chr12p13.31; q=0.007; occurring in 2 samples) [106,107], and *SPTLC3* (chr20p12.1 q=0.23; occurring in 2 samples) [108].

## DISCUSSION

In this study, we detected thousands of mutations in BCC samples, many of which were clustered in specific genes/regions or hotspots located in both coding and noncoding regions. Despite the small size of our dataset, our results are in line with those of previous genomic analyses of coding mutations in BCC [14,15], which confirms the reliability of our study. We believe that our results may give valuable insights related to general characteristics of mutations such as mutational burden or mutational signatures and in terms of genes identified as recurrently mutated in coding regions.

Moreover, we extended our analysis to noncoding parts of the genes, which altogether were responsible for ∼50% of the mutations identified by standard WES approach. Variants in such areas have usually been ignored in previous BCC genetic studies. Many of the identified noncoding hotspots were located in sequences of genes functionally related to cancer or more specifically to UV radiation-related skin cancers. Some of them were reported before in melanoma or identified by us in melanoma TCGA samples, the cancer type most intensively studied in terms of mutations in noncoding regions [109,110]. Below, we briefly describe the cancer-related role of the three most interesting genes with hotspot mutations in noncoding regions, i.e., *BAD*, *DHODH*, and *CHCHD2*. Interestingly, all these genes have functions related to mitochondrial activity.

Of all the hotspots detected in our study, the most frequently mutated was the hotspot located in the 3’UTR of *BAD*. This hotspot had several different mutations affecting 2 nucleotide positions (142 and 143 nt downstream of the stop codon). Due to these mutations, *BAD* was also classified as being highly mutated in the 3’UTR and as the top most significant potential cancer driver. Consistently, the hotspot and several other positions in the 3’UTR are frequently mutated in melanoma but not in other cancers. BAD belongs to the BCL-2 family, consisting of both proapoptotic and antiapoptotic proteins. It promotes cell death by inducing mitochondrial outer membrane permeabilization (MOMP), allowing the release of cytochrome c, and by antagonizing (dimerizing with) antiapoptotic BCL-2 proteins [111,112]. On the other hand, phosphorylated BAD may also have antiapoptotic properties, e.g., promoting the survival of melanocytes [113,114]. Other functions of BAD include regulation of mitochondrial metabolism (regulation of voltage-dependent anion channels and metabolite passage through the outer mitochondrial membrane) and dynamics (regulation of shape changes) [115–121]. Although BAD has not been previously implicated in skin cancers, loss or downregulation of other proapoptotic members of the BCL-2 family, i.e., BAX and PUMA, has been shown to promote the development of BCC, SCC, and cutaneous melanoma [122,123]. Therefore, a similar effect may be induced by mutations causing more efficient downregulation of BAD. *CHCHD2* is a gene with frequent mutations in the 5’UTR, the hotspot mutation c.-77C>T and the recurrent mutation c.-134G>A (77 and 134 upstream of the start codon). Based on the 5’UTR mutations, *CHCHD2* was classified as a high priority cancer driver. We showed that the *CHCHD2* 5’UTR (predominantly the hotspot position) was also frequently mutated (8%) in the SKCM TCGA samples, which also showed the additional recurrent mutation c.-74C>T. The 5’UTR mutations were also found in whole-genome sequenced Australian melanoma samples [56]. The role of the gene has not been intensively studied in cancer, but it was shown that under hypoxic conditions, CHCHD2 is translocated from the mitochondrial intermembrane space to the nucleus, where it binds an oxygen-responsive element in the promoter of *cytochrome oxidase 4I2* (*COX4I2*), encoding a subunit of complex IV of the electron transport chain, and increases its expression. Consequently, *CHCHD2* knockdown downregulates *COX4I2* and decreases cell oxygen consumption [124]. It was also shown that CHCHD2 is a negative regulator of mitochondria-mediated apoptosis [125]. Liu et al. showed that CHCHD2 interacts with antiapoptotic BCL-XL (from the BCL-2 family), which leads to inhibition of proapoptotic BAX and consequently decreases MOMP and apoptosis. In addition, it was shown that CHCHD2 dysregulates multiple genes that play a role in cell migration and cancer metastasis and that its expression is higher in cell lines derived from more aggressive breast tumors [126]. Consistent with the function of CHCHD2 related to mitochondrial metabolism, we found that all BCC/SKCM hotspot/recurrent mutations coincided with and impaired two distinct binding sites of GABP-alpha. As GABP-alpha is known to be a transcription factor involved in the regulation of cellular energy metabolism and cell cycle regulation [127], this finding might hint at a functional role of the mutations in cancer. Of note, germline missense mutations in *CHCHD2* are associated with autosomal dominant Parkinson’s disease [128].

*DHODH* is a gene that showed frequent mutations in the Kozak sequence of the 5’UTR, with hotspot mutations encompassing two different substitutions, c.-5G>A and c.-5_-4delinsAA (4 and 5 nt upstream of the start codon). Based on the identified mutations, *DHODH* was classified as a candidate cancer driver. The analysis of the entire TCGA cohort (∼10K samples from 33 cancer types) showed that no other cancer had mutations in the hotspot or the 5’UTR, indicating that the mutations were BCC-specific. Although *DHODH* 5’UTR mutations have never been reported before in any cancer, it was shown very recently that *DHODH* plays a key role in the carcinogenesis of SCC and other UV radiation-induced skin cancers and facilitates the development of precancerous skin lesions [54,55]. Hosseini et al. showed that the DHODH protein level and enzymatic activity are markedly upregulated in irradiated skin and that an increased level of DHODH sensitizes the skin to UV irradiation-induced damage. It was also shown that DHODH is upregulated in melanoma, in which DHODH inhibition leads to a marked decrease in tumor growth both *in vitro* and in mouse xenograft studies [129]. DHODH inactivation inhibits cell proliferation and induces cell cycle arrest at the S phase in BCL-2 (pro-apoptotic) deficient melanoma cells (Liu et al. 2017). DHODH is embedded in the inner mitochondrial membrane, and its canonical role is in the oxidation of dihydroorotate to orotate, an important step in *de novo* pyrimidine synthesis (which is important in replication and DNA repair). However, a side product of the pathway, ubiquinol (QH2), is a source of electrons in the electron transport chain, and DHODH also plays a role in alternative (glucose-independent) respiration (utilizing amino acids as an energy source) [54,55,130], facilitating cancer development in hypoxic conditions. In addition, it was found that in esophageal SCC, elevated DHODH levels promote cell proliferation by stabilizing β-catenin [131]. The functional effects of the mutations may result from alteration of the Kozak sequence but also the creation of an NFAT1 transcription factor binding site, which is not present in the wild-type sequence. NFAT1 is a widely distributed isoform of the NFAT family of transcription factors and is expressed in tumor cells and the tumor microenvironment [132]. The constitutive activation and overexpression of NFAT1 in many cancer types promote the transcription of genes that are crucial for cancer development and progression, including *COX2*, *MMP7*, *MMP9*, and *MDM2* [133,134].

It is worth noting that noncoding mutations in only the promoters of *TERT* and *DPH3* were previously analyzed in BCC [27,135,136]; *TERT* and *DPH3* are known to be mutated in many cancers, including melanoma [109,110]. However, as our WES design generally did not cover promoter regions, we did not look for or detect these mutations.

Additionally, the whole-genome CNA analysis allowed us to detect two highly significant chromosome-level CNAs. In addition to the expected deletion of chr9q, consistent with the loss of heterozygosity of *PTCH1*, we also detected frequent duplication/amplification of chr9p, encompassing the *PDL1* and *PDL2* genes (which encode the two immune checkpoint proteins PD-L1 and PD-L2, the overexpression of which enables cancer cells to evade the host immune system). Copy number gains of *PDL1* have been observed only in one case of metastatic BCC [95]. The patient, who was otherwise resistant to *vismodegib* and *sonidegib*, demonstrated a dramatic response to *nivolumab* (an anti-PD-1 antibody blocking the PD-1/PD-L1 interaction), which strongly suggested that the copy number gain may be a biomarker of sensitivity to anti-PD-1/PD-L1 checkpoint treatments [95]. It was also shown in an independent study that some patients (up to ∼40%) with advanced BCC (not tested for *PDL1* amplification) respond to *pembrolizumab* (another anti-PD-1 antibody) [95,137]. Therefore, assessment of copy number gains of the *PDL1/PDL2* region may help to rationalize such treatment; however, further study with a larger group of samples is required.

Finally, we would like to note the apparent limitations of the study. As it was intended to be a preliminary evaluation of noncoding mutations in BCC, we analyzed only a small number of samples, and as such, we limited the characterization of the identified variants to computational analyses.

In summary, in this study utilizing WES BCC data, we revealed not only mutations in coding regions of previously known BCC-related genes but also frequent mutations in noncoding regions of cancer-related genes, some of which may be strong candidates for new BCC drivers. Although the functional role of the individual identified genes/mutations requires further experimental interrogations, our results provide a strong basis for further analyses of noncoding variants in BCC and other cancer types.

## MATERIALS AND METHODS

### Sample collection and DNA preparation

A total of 27 pairs of tissue (tumor and normal adjacent healthy skin) were collected from the Department of Plastic Surgery, St. Josef Hospital, Catholic Clinics of the Ruhr Peninsula, Essen, Germany. While excising the BCC tissues with cold steel under local anesthesia, 4-mm punch biopsies were taken from the center of the tumor and from nonlesional epithelial skin (as normal, intraindividual controls). These samples were immediately placed in RNAlater (Qiagen, Hilden, Germany) and stored at −80 °C. Tissue homogenization was performed with stainless steel beads of 5 mm (Qiagen) and TissueLyser LT (Qiagen). DNA was extracted with an AllPrep DNA/RNA/miRNA Universal Kit (Qiagen) according to the manufacturer’s protocol. All samples were quantified using a NanoDrop One (Thermo Scientific, Waltham, USA) and Qubit fluorometer 3.0 (Invitrogen) (Qubit dsDNA HS Assay (Life Technologies, Carlsbad, USA)), and DNA size and quality were tested using gel electrophoresis.

### Exome sequencing and data processing

The library was prepared with 200 ng of high-quality DNA using the SureSelectXT Library Prep Kit (Agilent). A SureSelectXT Human All Exon V6 kit (Agilent) was used for exome capture. Sequencing was performed on an Illumina NovaSeq 6000 (San Diego, USA), generating 2x 100 bp paired-end reads. Library preparation, exome enrichment, and sequencing were performed at CeGaT, Tuebingen, Germany. Demultiplexing of the sequencing reads was performed with Illumina bcl2fastq (2.19). Adapters were trimmed with Skewer (version 0.2.2) [138]. The Phred score was given with Illumina standard Phred encoding (offset +33). For each sample, two FASTQ files corresponding to forward and reverse reads were obtained. Paired-end reads were aligned to hg38 using BWA. PCR duplicates were marked and removed with the Picard package. Indel realignments with known sites and base quality score recalibration were performed with GATK version 4.1.2.0. SAM to BAM conversion was done using SAMtools. Somatic single-nucleotide variants were called with MuTect2 (version 4.1.0.0. with the use of the tumor-normal mode). To distinguish somatic from germline mutations, we relied on the gnomAD database (version 2.1.1) as a public germline mutation source. We also generated a panel of normals (PON) file, which contained a set of variants that were detected by MuTect2 when run on a set of normal samples (but these were not germline mutations). From the VCF (variant call format) files, we extracted somatic mutation calls with PASS and panel of normals annotation. We also added information about the localization of mutations in gene subregions (CDS, 5’UTR, 3’UTR, or introns) by use of an in-house Python script. From the list of somatic mutations, we additionally removed those that did not fulfill the following criteria: (i) at least five alternative allele-supporting reads in a tumor sample; (ii) frequency of alternative allele-supporting reads in a tumor sample of at least 0.05; and (iii) frequency of alternative allele-supporting reads in the tumor sample at least 5× higher than that in the corresponding normal sample.

### Validation of mutations

A panel of 51 mutations detected by WES was validated by Sanger sequencing of the appropriate PCR fragments amplified with primers shown in S8 Table. All fragments were sequenced in two directions with the BigDye v3.1 kit (Applied Biosystems, Foster City, CA, USA), and the sequencing reactions were separated with capillary electrophoresis (POP7 polymer; ABI Prism 3130xl apparatus; Applied Biosystems, Foster City, CA, USA) according to the standard manufacturer’s recommendations.

### Mutational signature analysis

To analyze mutational signatures, we used the web application Mutational Signatures in Cancer (MuSiCa; http://bioinfo.ciberehd.org/GPtoCRC/en/tools.html [139]), allowing the visualization of the somatic mutational profile of each analyzed sample and estimation of the contribution values of the predefined mutational signatures ([140]; Catalogue Of Somatic Mutations In Cancer, COSMIC 2020). Samples BCC14 and BCC21 were excluded from the signature analysis due to an insufficient number of mutations.

### Identification of hotspots, frequently mutated genes, and cancer drivers

We defined genomic positions mutated in at least 3 samples as hotspots. Mutations occurring in directly adjacent nucleotides were merged into one hotspot.

We defined genes with nonsynonymous mutations in a coding region in at least 5 samples, with mutations in a 5’UTR, in at least 4 samples, with mutations in a 3’UTR in at least 4 samples, and with mutations in introns (up to 40 nt from exon/intron boundaries) in at least 5 samples as frequently mutated. From the analysis, we excluded genes known to be commonly hypermutated with passenger mutations as a result of the increased background mutation rate but not related to cancer, listed in [141]. To distinguish synonymous from nonsynonymous mutations, we used the SnpEff - genetic variant annotation and functional effect prediction toolbox [142], available on the Subio platform (Subio, Inc., Kagoshima, Japan, http://www.subio.jp). We also considered splice-site mutations located in introns up to +/-2 nt from exons as coding region mutations.

OncodriveFML [83] was run using the CADD score (hg38, version 1.6). The signature method was set as a complement, the statistical method was set to “amean”, and indels were included in the analysis using a max method (max_consecutive was set to 7 as default).

### Copy number analysis

To identify chromosome arm-level and focal regions that were significantly amplified or deleted, we used GISTIC2 [94] with the following parameters: threshold for copy number amplifications and deletions, 0.2; confidence level to calculate the region containing a driver, 0.9; broad-level analysis; and the arm peel method to reduce noise.

To validate CNAs involving chromosome 9, i.e., chr9p duplications/amplifications (affecting *JAK2*, *PDL1*/*CD274*, and *PDL2*/*CD273*) and chr9q deletions (affecting *PTCH1*), we designed and generated an MLPA assay covering the entire chromosome 9. In total, the assay consisted of 20 probes, including (i) 7 probes distributed over the chr9p (n=5) and chr9q (n=2) arms, 2 probes located in or in close proximity to *JAK2*, *PDL2*, *PDL1*, and *PTCH1* (in total 8 gene-specific probes), and 5 control probes (located on different chromosomes outside of chromosome 9 and regions of known cancer-related genes). The sequences and detailed characteristics of all probes as well as their exact positions are shown in S9 Table and are schematically depicted in Fig 6C.

The MLPA probes and the probe-set layout were designed according to a previously proposed and well-validated strategy [143,144].

### TCGA analysis

To compare the mutations recurring in BCC with mutations in other cancers, we used WES-generated somatic mutation datasets of 10,369 samples representing 33 cancer types generated and deposited in the TCGA repository (http://cancergenome.nih.gov). The full names and abbreviations of all TCGA cancer types are shown in S4 Table. Somatic mutations were identified against matched normal samples with the use of the standard TCGA pipeline (including the Mutect2, Muse, Varscan, and SomaticSnipper algorithms). We extracted somatic mutation calls (with PASS annotation only) localized in the annotated exons of *BAD*, *DHODH*, *CHCHD2*, *FLG*, and *FLG2* (exon sequences were extended by 2 nt to enable identification of intronic splice-site mutations). The extraction was performed as described in our earlier study [145] with a set of in-house Python scripts available at (https://github.com/martynaut/mirnaome_somatic_mutations).

### Mutations visualization

All mutations were annotated according to HGVS nomenclature (at the transcript and protein levels), and the effects of mutations were defined using the Ensembl Variant Effect Predictor (VEP) tool. For visualization of mutations on gene maps, we used ProteinPaint from St. Jude Children’s Research Hospital – PeCan Data Portal [146]. The protein domains visualized on gene maps were positioned according to UniProt data [147]. The comutation plot showing frequently mutated genes was created with the use of the Python library CoMut [148].

### Analysis of RNA regulatory motifs

Target predictions were performed with the TargetScan Custom (release 5.2) web tool [149]. The secondary RNA structures were predicted using mfold software [150] with default parameters. RNA sequence/structure functional motifs and transcription factor binding sites were analyzed with the RegRNA 2.0 [151] and MotifMap [65] web tools.

### Statistics

Specific statistical tests are indicated in the text, and a *p*-value <0.05 was considered significant. If necessary, *p*-values were corrected for multiple tests with the Benjamini-Hochberg procedure.

## Author Contributions

Conceptualization: Michael Sand, Piotr Kozlowski

Data curation: Paulina Nawrocka, Paulina Galka-Marciniak, Martyna Urbanek-Trzeciak

Formal analysis: Paulina Nawrocka, Paulina Galka-Marciniak, Martyna Urbanek-Trzeciak, Natalia Szostak, Piotr Kozlowski

Funding acquisition: Piotr Kozlowski

Investigation: Paulina Nawrocka, Paulina Galka-Marciniak, Martyna Urbanek-Trzeciak, Ilamathi M., Piotr Kozlowski

Methodology: Paulina Nawrocka, Paulina Galka-Marciniak, Martyna Urbanek-Trzeciak, Piotr Kozlowski

Project administration: Paulina Nawrocka, Piotr

Kozlowski Resources: Laura Susok, Michael Sand

Software: Paulina Nawrocka, Paulina Galka-Marciniak, Martyna Urbanek-Trzeciak, Natalia Szostak, Anna Philips

Supervision: Piotr Kozlowski

Validation: Paulina Nawrocka

Visualization: Paulina Nawrocka, Paulina Galka-Marciniak, Piotr Kozlowski

Writing - original draft: Paulina Nawrocka, Piotr Kozlowski

Writing - review & editing: Paulina Nawrocka, Paulina Galka-Marciniak, Martyna Urbanek-Trzeciak, Ilamathi M., Natalia Szostak, Anna Philips, Laura Susok, Michael Sand, Piotr Kozlowski

## Supporting information

**S1 Fig.**
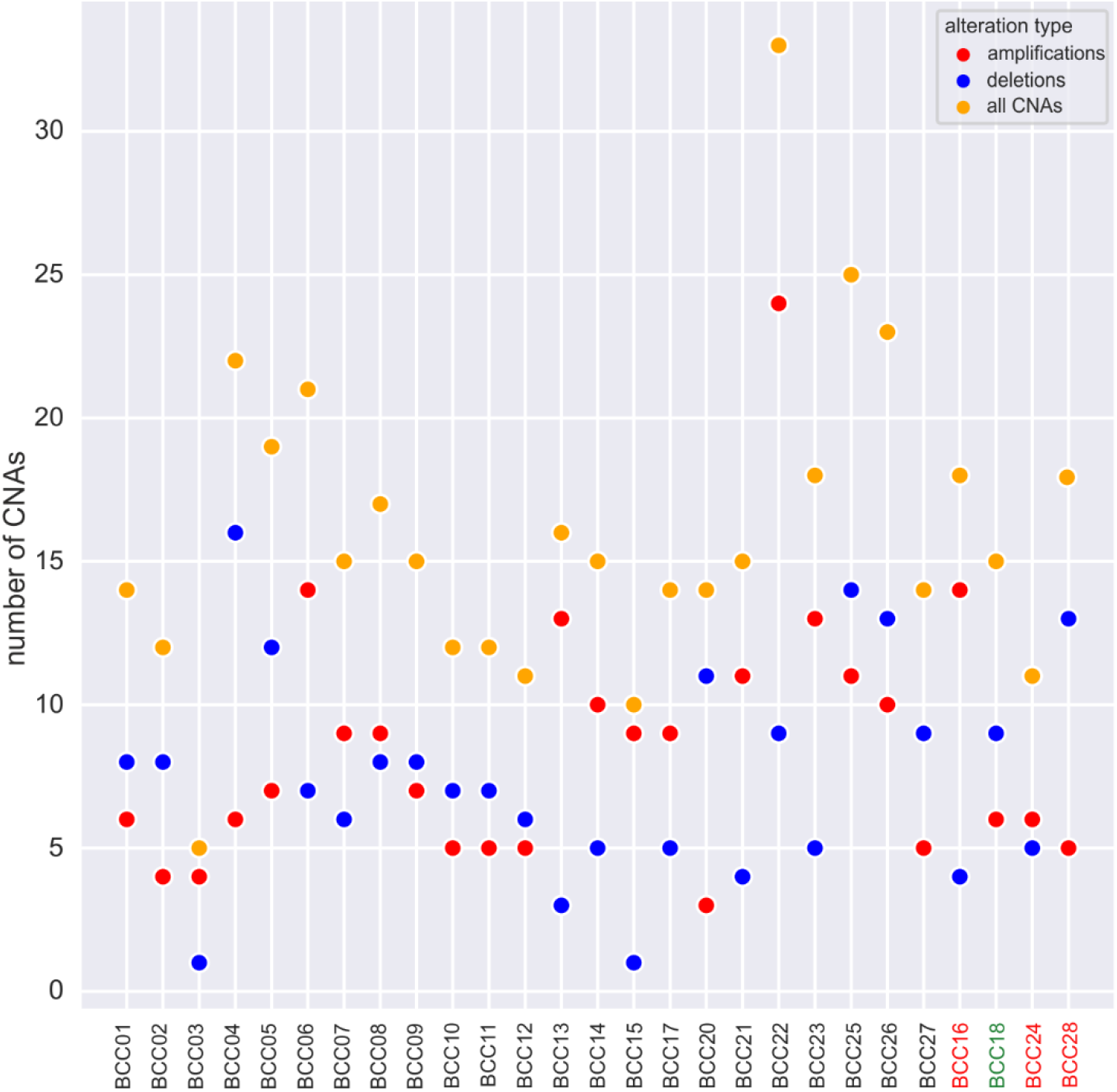
Concurrence of KNSTRN mutations and CNAs. The number of amplifications (red dots), deletions (blue dots), and total CNAs (orange dots) in samples with and without the *KNSTRN* mutations (red font - hotspot mutation; green font - other mutation).

**S2 Fig.**
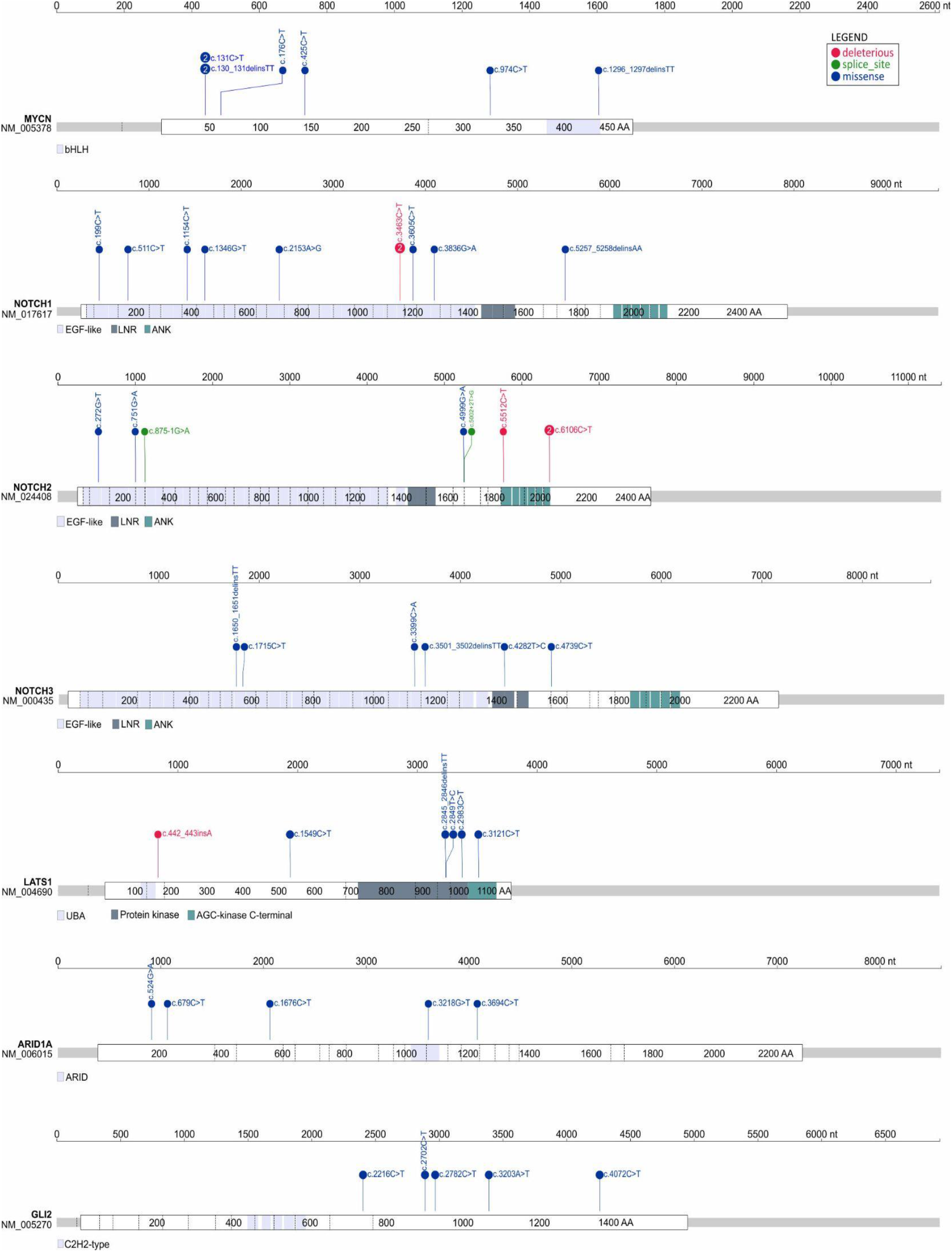
Distribution of the identified mutations in the selected genes frequently mutated in coding sequence. A)-F) Maps of *MYCN*, *NOTCH1*, *NOTCH2*, *NOTCH3*, *LATS1,* and *ARID1A*, respectively.

**S3 Fig.**
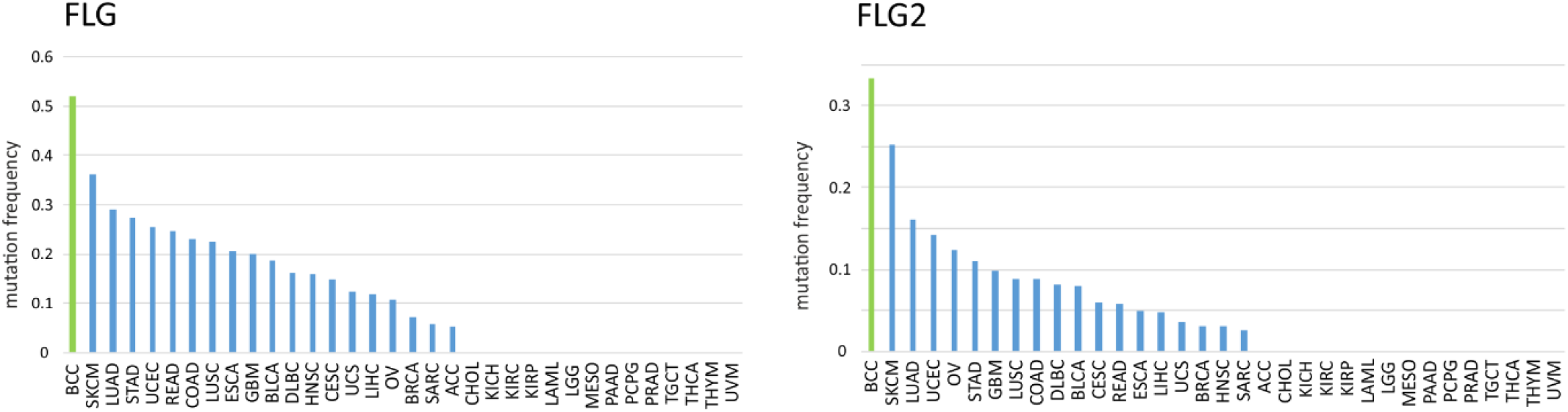
Frequency of mutations in the *FLG* and *FLG2* genes in BCC and other cancer types. Bar graphs showing the frequency of mutations in A) *FLG*, and B) *FLG2* in BCC (current study, green bar) and TCGA cancer types.

**S4 Fig.**
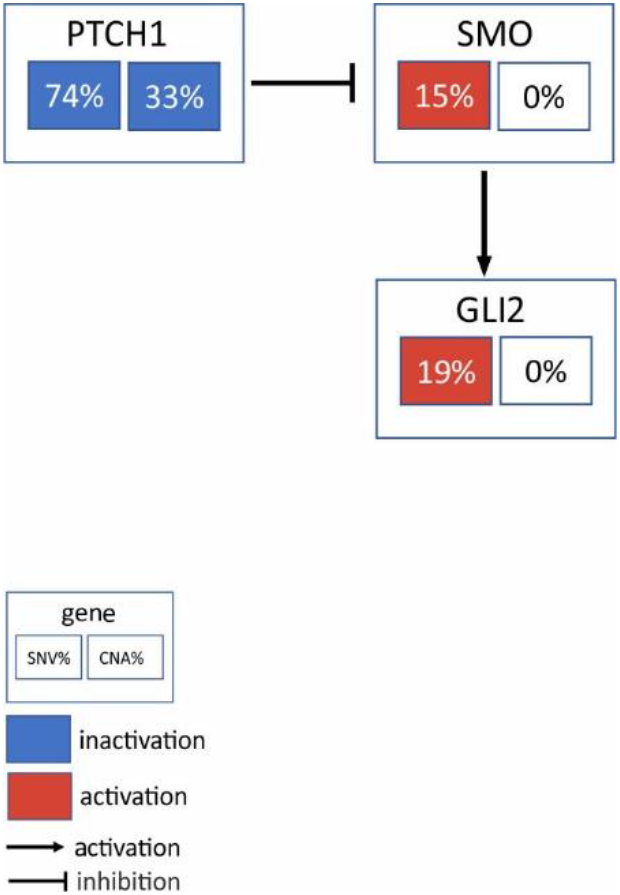
Frequency of activating (red) and deactivating (blue) mutations in genes of the hedgehog pathway.

S1 Table. List of somatic mutations in BCC.

(XLSX)

S2 Table. List of somatic mutations validated by Sanger sequencing.

(XLSX)

S3 Table. List of hotspot mutations.

(XLSX)

S4 Table. TCGA abbreviations and full cancer/project names.

(XLSX)

S5 Table. List of frequently mutated genes.

(XLSX)

S6 Table. List of driver genes identified by the OncodriveFML analysis.

(XLSX)

S7 Table. List of significant focal CNAs.

(XLSX)

S8 Table. List of the primers used for PCR.

(XLSX)

S9 Table. List of the probes used for the MLPA assay.

(XLSX)

## REFERENCES

1. Samarasinghe V, Madan V, Lear JT. Focus on Basal cell carcinoma. J Skin Cancer. 2011;2011: 328615. doi:10.1155/2011/328615

2. Puig S, Berrocal A. Management of high-risk and advanced basal cell carcinoma. Clin Transl Oncol Off Publ Fed Span Oncol Soc Natl Cancer Inst Mex. 2015;17: 497–503. doi:10.1007/s12094-014-1272-9

3. Pellegrini C, Maturo MG, Di Nardo L, Ciciarelli V, Gutiérrez García-Rodrigo C, Fargnoli MC. Understanding the Molecular Genetics of Basal Cell Carcinoma. Int J Mol Sci. 2017;18. doi:10.3390/ijms18112485

4. Evans DG, Howard E, Giblin C, Clancy T, Spencer H, Huson SM, et al. Birth incidence and prevalence of tumor-prone syndromes: estimates from a UK family genetic register service. Am J Med Genet A. 2010;152A: 327–332. doi:10.1002/ajmg.a.33139

5. Sexton M, Jones DB, Maloney ME. Histologic pattern analysis of basal cell carcinoma. Study of a series of 1039 consecutive neoplasms. J Am Acad Dermatol. 1990;23: 1118–1126. doi:10.1016/0190-9622(90)70344-h

6. Rippey JJ. Why classify basal cell carcinomas? Histopathology. 1998;32: 393–398. doi:10.1046/j.1365-2559.1998.00431.x

7. Raasch BA, Buettner PG, Garbe C. Basal cell carcinoma: histological classification and body-site distribution. Br J Dermatol. 2006;155: 401–407. doi:10.1111/j.1365-2133.2006.07234.x

8. Lo JS, Snow SN, Reizner GT, Mohs FE, Larson PO, Hruza GJ. Metastatic basal cell carcinoma: report of twelve cases with a review of the literature. J Am Acad Dermatol. 1991;24: 715–719. doi:10.1016/0190-9622(91)70108-e

9. Ting PT, Kasper R, Arlette JP. Metastatic basal cell carcinoma: report of two cases and literature review. J Cutan Med Surg. 2005;9: 10–15. doi:10.1007/s10227-005-0027-1

10. Crowson AN, Magro CM, Kadin ME, Stranc M. Differential expression of the bcl-2 oncogene in human basal cell carcinoma. Hum Pathol. 1996;27: 355–359. doi:10.1016/s0046-8177(96)90108-2

11. Crowson AN. Basal cell carcinoma: biology, morphology and clinical implications. Mod Pathol Off J U S Can Acad Pathol Inc. 2006;19 Suppl 2: S127–147. doi:10.1038/modpathol.3800512

12. Epstein EH. Basal cell carcinomas: attack of the hedgehog. Nat Rev Cancer. 2008;8: 743–754. doi:10.1038/nrc2503

13. Iwasaki JK, Srivastava D, Moy RL, Lin HJ, Kouba DJ. The molecular genetics underlying basal cell carcinoma pathogenesis and links to targeted therapeutics. J Am Acad Dermatol. 2012;66: e167–178. doi:10.1016/j.jaad.2010.06.054

14. Jayaraman SS, Rayhan DJ, Hazany S, Kolodney MS. Mutational Landscape of Basal Cell Carcinomas by Whole-Exome Sequencing. J Invest Dermatol. 2014;134: 213–220. doi:10.1038/jid.2013.276

15. Bonilla X, Parmentier L, King B, Bezrukov F, Kaya G, Zoete V, et al. Genomic analysis identifies new drivers and progression pathways in skin basal cell carcinoma. Nat Genet. 2016;48: 398–406. doi:10.1038/ng.3525

16. Verkouteren J a. C, Ramdas KHR, Wakkee M, Nijsten T. Epidemiology of basal cell carcinoma: scholarly review. Br J Dermatol. 2017;177: 359–372. doi:10.1111/bjd.15321

17. Peris K, Fargnoli MC, Garbe C, Kaufmann R, Bastholt L, Seguin NB, et al. Diagnosis and treatment of basal cell carcinoma: European consensus-based interdisciplinary guidelines. Eur J Cancer Oxf Engl 1990. 2019;118: 10–34. doi:10.1016/j.ejca.2019.06.003

18. Gailani MR, Leffell DJ, Ziegler A, Gross EG, Brash DE, Bale AE. Relationship between sunlight exposure and a key genetic alteration in basal cell carcinoma. J Natl Cancer Inst. 1996;88: 349–354. doi:10.1093/jnci/88.6.349

19. Maloney ME. Arsenic in Dermatology. Dermatol Surg Off Publ Am Soc Dermatol Surg Al. 1996;22: 301–304. doi:10.1111/j.1524-4725.1996.tb00322.x

20. Rubin AI, Chen EH, Ratner D. Basal-cell carcinoma. N Engl J Med. 2005;353: 2262–2269. doi:10.1056/NEJMra044151

21. Palacios-Álvarez I, González-Sarmiento R, Fernández-López E. Gorlin Syndrome. Actas Dermo-Sifiliográficas Engl Ed. 2018;109: 207–217. doi:10.1016/j.adengl.2018.02.002

22. Diederichs S, Bartsch L, Berkmann JC, Fröse K, Heitmann J, Hoppe C, et al. The dark matter of the cancer genome: aberrations in regulatory elements, untranslated regions, splice sites, non-coding RNA and synonymous mutations. EMBO Mol Med. 2016;8: 442–457. doi:10.15252/emmm.201506055

23. Leppek K, Das R, Barna M. Functional 5’ UTR mRNA structures in eukaryotic translation regulation and how to find them. Nat Rev Mol Cell Biol. 2018;19: 158–174. doi:10.1038/nrm.2017.103

24. Lord J, Gallone G, Short PJ, McRae JF, Ironfield H, Wynn EH, et al. Pathogenicity and selective constraint on variation near splice sites. Genome Res. 2019;29: 159–170. doi:10.1101/gr.238444.118

25. Horn S, Figl A, Rachakonda PS, Fischer C, Sucker A, Gast A, et al. TERT promoter mutations in familial and sporadic melanoma. Science. 2013;339: 959–961. doi:10.1126/science.1230062

26. Huang FW, Hodis E, Xu MJ, Kryukov GV, Chin L, Garraway LA. Highly recurrent TERT promoter mutations in human melanoma. Science. 2013;339: 957–959. doi:10.1126/science.1229259

27. Maturo MG, Rachakonda S, Heidenreich B, Pellegrini C, Srinivas N, Requena C, et al. Coding and noncoding somatic mutations in candidate genes in basal cell carcinoma. Sci Rep. 2020;10: 8005. doi:10.1038/s41598-020-65057-2

28. Urbanek-Trzeciak MO, Galka-Marciniak P, Nawrocka PM, Kowal E, Szwec S, Giefing M, et al. Pan-cancer analysis of somatic mutations in miRNA genes. EBioMedicine. 2020;61: 103051. doi:10.1016/j.ebiom.2020.103051

29. Sand M, Skrygan M, Sand D, Georgas D, Hahn SA, Gambichler T, et al. Expression of microRNAs in basal cell carcinoma. Br J Dermatol. 2012;167: 847–855. doi:10.1111/j.1365-2133.2012.11022.x

30. Sand M, Hessam S, Amur S, Skrygan M, Bromba M, Stockfleth E, et al. Expression of oncogenic miR-17-92 and tumor suppressive miR-143-145 clusters in basal cell carcinoma and cutaneous squamous cell carcinoma. J Dermatol Sci. 2017;86: 142–148. doi:10.1016/j.jdermsci.2017.01.012

31. Sand M, Bromba A, Sand D, Gambichler T, Hessam S, Becker JC, et al. Dicer Sequencing, Whole Genome Methylation Profiling, mRNA and smallRNA Sequencing Analysis in Basal Cell Carcinoma. Cell Physiol Biochem Int J Exp Cell Physiol Biochem Pharmacol. 2019;53: 760–773. doi:10.33594/000000171

32. Rheinbay E, Nielsen MM, Abascal F, Wala JA, Shapira O, Tiao G, et al. Analyses of non-coding somatic drivers in 2,658 cancer whole genomes. Nature. 2020;578: 102–111. doi:10.1038/s41586-020-1965-x

33. Sakthikumar S, Roy A, Haseeb L, Pettersson ME, Sundström E, Marinescu VD, et al. Whole-genome sequencing of glioblastoma reveals enrichment of non-coding constraint mutations in known and novel genes. Genome Biol. 2020;21: 127. doi:10.1186/s13059-020-02035-x

34. Lawrence MS, Stojanov P, Mermel CH, Garraway LA, Golub TR, Meyerson M, et al. Discovery and saturation analysis of cancer genes across 21 tumor types. Nature. 2014;505: 495–501. doi:10.1038/nature12912

35. Supek F, Miñana B, Valcárcel J, Gabaldón T, Lehner B. Synonymous Mutations Frequently Act as Driver Mutations in Human Cancers. Cell. 2014;156: 1324–1335. doi:10.1016/j.cell.2014.01.051

36. Bin Y, Wang X, Zhao L, Wen P, Xia J. An analysis of mutational signatures of synonymous mutations across 15 cancer types. BMC Med Genet. 2019;20: 190. doi:10.1186/s12881-019-0926-4

37. Sharma Y, Miladi M, Dukare S, Boulay K, Caudron-Herger M, Groß M, et al. A pan-cancer analysis of synonymous mutations. Nat Commun. 2019;10: 2569. doi:10.1038/s41467-019-10489-2

38. Hase K, Kimura S, Takatsu H, Ohmae M, Kawano S, Kitamura H, et al. M-Sec promotes membrane nanotube formation by interacting with Ral and the exocyst complex. Nat Cell Biol. 2009;11: 1427–1432. doi:10.1038/ncb1990

39. Jia L, Zhou Z, Liang H, Wu J, Shi P, Li F, et al. KLF5 promotes breast cancer proliferation, migration and invasion in part by upregulating the transcription of TNFAIP2. Oncogene. 2016;35: 2040–2051. doi:10.1038/onc.2015.263

40. Xie Y, Wang B. Downregulation of TNFAIP2 suppresses proliferation and metastasis in esophageal squamous cell carcinoma through activation of the Wnt/β-catenin signaling pathway. Oncol Rep. 2017;37: 2920–2928. doi:10.3892/or.2017.5557

41. Jia L, Shi Y, Wen Y, Li W, Feng J, Chen C. The roles of TNFAIP2 in cancers and infectious diseases. J Cell Mol Med. 2018;22: 5188–5195. doi:10.1111/jcmm.13822

42. Kodach LL, Jacobs RJ, Heijmans J, van Noesel CJM, Langers AMJ, Verspaget HW, et al. The role of EZH2 and DNA methylation in the silencing of the tumour suppressor RUNX3 in colorectal cancer. Carcinogenesis. 2010;31: 1567–1575. doi:10.1093/carcin/bgq147

43. Xie Q, Wang H, Heilman ER, Walsh MG, Haseeb MA, Gupta R. Increased expression of enhancer of Zeste Homolog 2 (EZH2) differentiates squamous cell carcinoma from normal skin and actinic keratosis. Eur J Dermatol EJD. 2014;24: 41–45. doi:10.1684/ejd.2013.2219

44. Rao RC, Chan MP, Andrews CA, Kahana A. EZH2, Proliferation Rate, and Aggressive Tumor Subtypes in Cutaneous Basal Cell Carcinoma. JAMA Oncol. 2016;2: 962–963. doi:10.1001/jamaoncol.2016.0021

45. Bödör C, Grossmann V, Popov N, Okosun J, O’Riain C, Tan K, et al. EZH2 mutations are frequent and represent an early event in follicular lymphoma. Blood. 2013;122: 3165–3168. doi:10.1182/blood-2013-04-496893

46. Donaldson-Collier MC, Sungalee S, Zufferey M, Tavernari D, Katanayeva N, Battistello E, et al. EZH2 oncogenic mutations drive epigenetic, transcriptional, and structural changes within chromatin domains. Nat Genet. 2019;51: 517–528. doi:10.1038/s41588-018-0338-y

47. Weaver TM, Liu J, Connelly KE, Coble C, Varzavand K, Dykhuizen EC, et al. The EZH2 SANT1 domain is a histone reader providing sensitivity to the modification state of the H4 tail. Sci Rep. 2019;9: 987. doi:10.1038/s41598-018-37699-w

48. Fang L, Seki A, Fang G. SKAP associates with kinetochores and promotes the metaphase-to-anaphase transition. Cell Cycle. 2009;8: 2819–2827. doi:10.4161/cc.8.17.9514

49. Lee CS, Bhaduri A, Mah A, Johnson WL, Ungewickell A, Aros CJ, et al. Recurrent point mutations in the kinetochore gene KNSTRN in cutaneous squamous cell carcinoma. Nat Genet. 2014;46: 1060–1062. doi:10.1038/ng.3091

50. Jaju PD, Nguyen CB, Mah AM, Atwood SX, Li J, Zia A, et al. Mutations in the Kinetochore Gene KNSTRN in Basal Cell Carcinoma. J Invest Dermatol. 2015;135: 3197–3200. doi:10.1038/jid.2015.339

51. Lu S, Wang R, Cai C, Liang J, Xu L, Miao S, et al. Small kinetochore associated protein (SKAP) promotes UV-induced cell apoptosis through negatively regulating pre-mRNA processing factor 19 (Prp19). PloS One. 2014;9: e92712. doi:10.1371/journal.pone.0092712

52. Marchion DC, Cottrill HM, Xiong Y, Chen N, Bicaku E, Fulp WJ, et al. BAD Phosphorylation Determines Ovarian Cancer Chemosensitivity and Patient Survival. Clin Cancer Res. 2011;17: 6356–6366. doi:10.1158/1078-0432.CCR-11-0735

53. Stickles XB, Marchion DC, Bicaku E, Al Sawah E, Abbasi F, Xiong Y, et al. BAD-mediated apoptotic pathway is associated with human cancer development. Int J Mol Med. 2015;35: 1081–1087. doi:10.3892/ijmm.2015.2091

54. Hosseini M, Dousset L, Mahfouf W, Serrano-Sanchez M, Redonnet-Vernhet I, Mesli S, et al. Energy Metabolism Rewiring Precedes UVB-Induced Primary Skin Tumor Formation. Cell Rep. 2018;23: 3621–3634. doi:10.1016/j.celrep.2018.05.060

55. Hosseini M, Dousset L, Michon P, Mahfouf W, Muzotte E, Bergeron V, et al. UVB-induced DHODH upregulation, which is driven by STAT3, is a promising target for chemoprevention and combination therapy of photocarcinogenesis. Oncogenesis. 2019;8: 52. doi:10.1038/s41389-019-0161-z

56. Hayward NK, Wilmott JS, Waddell N, Johansson PA, Field MA, Nones K, et al. Whole-genome landscapes of major melanoma subtypes. Nature. 2017;545: 175–180. doi:10.1038/nature22071

57. Dutton-Regester K, Gartner JJ, Emmanuel R, Qutob N, Davies MA, Gershenwald JE, et al. A highly recurrent RPS27 5’UTR mutation in melanoma. Oncotarget. 2014;5: 2912–2917.

58. Floristán A, Morales L, Hanniford D, Martinez C, Castellano-Sanz E, Dolgalev I, et al. Functional analysis of RPS27 mutations and expression in melanoma. Pigment Cell Melanoma Res. 2020;33: 466–479. doi:10.1111/pcmr.12841

59. Huang H-Y, Lin Y-C-D, Li J, Huang K-Y, Shrestha S, Hong H-C, et al. miRTarBase 2020: updates to the experimentally validated microRNA-target interaction database. Nucleic Acids Res. 2020;48: D148–D154. doi:10.1093/nar/gkz896

60. Salamov AA, Nishikawa T, Swindells MB. Assessing protein coding region integrity in cDNA sequencing projects. Bioinforma Oxf Engl. 1998;14: 384–390. doi:10.1093/bioinformatics/14.5.384

61. Peitsch MC. Large scale protein modelling and model repository. Proc Int Conf Intell Syst Mol Biol. 1997;5: 234–236.

62. Chang T-H, Huang H-Y, Hsu JB-K, Weng S-L, Horng J-T, Huang H-D. An enhanced computational platform for investigating the roles of regulatory RNA and for identifying functional RNA motifs. BMC Bioinformatics. 2013;14: S4. doi:10.1186/1471-2105-14-S2-S4

63. Shoshan E, Braeuer RR, Kamiya T, Mobley AK, Huang L, Vasquez ME, et al. NFAT1 Directly Regulates IL8 and MMP3 to Promote Melanoma Tumor Growth and Metastasis. Cancer Res. 2016;76: 3145–3155. doi:10.1158/0008-5472.CAN-15-2511

64. Shen Y, Song Z, Lu X, Ma Z, Lu C, Zhang B, et al. Fas signaling-mediated T H 9 cell differentiation favors bowel inflammation and antitumor functions. Nat Commun. 2019;10: 2924. doi:10.1038/s41467-019-10889-4

65. Daily K, Patel VR, Rigor P, Xie X, Baldi P. MotifMap: integrative genome-wide maps of regulatory motif sites for model species. BMC Bioinformatics. 2011;12: 495. doi:10.1186/1471-2105-12-495

66. Wang NJ, Sanborn Z, Arnett KL, Bayston LJ, Liao W, Proby CM, et al. Loss-of-function mutations in Notch receptors in cutaneous and lung squamous cell carcinoma. Proc Natl Acad Sci. 2011;108: 17761–17766. doi:10.1073/pnas.1114669108

67. Purdie KJ, Lambert SR, Teh M-T, Chaplin T, Molloy G, Raghavan M, et al. Allelic Imbalances and Microdeletions Affecting the PTPRD Gene in Cutaneous Squamous Cell Carcinomas Detected Using Single Nucleotide Polymorphism Microarray Analysis. Genes Chromosomes Cancer. 2007;46: 661–669. doi:10.1002/gcc.20447

68. Veeriah S, Brennan C, Meng S, Singh B, Fagin JA, Solit DB, et al. The tyrosine phosphatase PTPRD is a tumor suppressor that is frequently inactivated and mutated in glioblastoma and other human cancers. Proc Natl Acad Sci. 2009;106: 9435–9440. doi:10.1073/pnas.0900571106

69. Walia V, Prickett TD, Kim J-S, Gartner JJ, Lin JC, Zhou M, et al. Mutational and Functional Analysis of the Tumor-Suppressor PTPRD in Human Melanoma. Hum Mutat. 2014;35: 1301–1310. doi:10.1002/humu.22630

70. Peyser ND, Du Y, Li H, Lui V, Xiao X, Chan TA, et al. Loss-of-Function PTPRD Mutations Lead to Increased STAT3 Activation and Sensitivity to STAT3 Inhibition in Head and Neck Cancer. PLOS ONE. 2015;10: e0135750. doi:10.1371/journal.pone.0135750

71. Szaumkessel M, Wojciechowska S, Janiszewska J, Zemke N, Byzia E, Kiwerska K, et al. Recurrent epigenetic silencing of the PTPRD tumor suppressor in laryngeal squamous cell carcinoma. Tumor Biol. 2017;39: 1010428317691427. doi:10.1177/1010428317691427

72. Adib E, Klonowska K, Giannikou K, Do KT, Pruitt-Thompson S, Bhushan K, et al. Phase II Clinical Trial of Everolimus in a Pan-Cancer Cohort of Patients with mTOR Pathway Alterations. Clin Cancer Res. 2021 [cited 27 May 2021]. doi:10.1158/1078-0432.CCR-20-4548

73. Wu Z, Hansmann B, Meyer-Hoffert U, Gläser R, Schröder J-M. Molecular Identification and Expression Analysis of Filaggrin-2, a Member of the S100 Fused-Type Protein Family. PLoS ONE. 2009;4. doi:10.1371/journal.pone.0005227

74. Leman G, Moosbrugger-Martinz V, Blunder S, Pavel P, Dubrac S. 3D-Organotypic Cultures to Unravel Molecular and Cellular Abnormalities in Atopic Dermatitis and Ichthyosis Vulgaris. Cells. 2019;8. doi:10.3390/cells8050489

75. Liu H, Toman RE, Goparaju SK, Maceyka M, Nava VE, Sankala H, et al. Sphingosine kinase type 2 is a putative BH3-only protein that induces apoptosis. J Biol Chem. 2003;278: 40330–40336. doi:10.1074/jbc.M304455200

76. Igarashi N, Okada T, Hayashi S, Fujita T, Jahangeer S, Nakamura S. Sphingosine kinase 2 is a nuclear protein and inhibits DNA synthesis. J Biol Chem. 2003;278: 46832–46839. doi:10.1074/jbc.M306577200

77. Maceyka M, Sankala H, Hait NC, Le Stunff H, Liu H, Toman R, et al. SphK1 and SphK2, sphingosine kinase isoenzymes with opposing functions in sphingolipid metabolism. J Biol Chem. 2005;280: 37118–37129. doi:10.1074/jbc.M502207200

78. Van Brocklyn JR, Jackson CA, Pearl DK, Kotur MS, Snyder PJ, Prior TW. Sphingosine kinase-1 expression correlates with poor survival of patients with glioblastoma multiforme: roles of sphingosine kinase isoforms in growth of glioblastoma cell lines. J Neuropathol Exp Neurol. 2005;64: 695–705. doi:10.1097/01.jnen.0000175329.59092.2c

79. Gao P, Smith CD. Ablation of sphingosine kinase-2 inhibits tumor cell proliferation and migration. Mol Cancer Res MCR. 2011;9: 1509–1519. doi:10.1158/1541-7786.MCR-11-0336

80. Su X, Malouf GG, Chen Y, Zhang J, Yao H, Valero V, et al. Comprehensive analysis of long non-coding RNAs in human breast cancer clinical subtypes. Oncotarget. 2014;5: 9864–9876.

81. Segers VFM, Dugaucquier L, Feyen E, Shakeri H, De Keulenaer GW. The role of ErbB4 in cancer. Cell Oncol Dordr. 2020;43: 335–352. doi:10.1007/s13402-020-00499-4

82. Sand M, Gambichler T, Skrygan M, Sand D, Scola N, Altmeyer P, et al. Expression levels of the microRNA processing enzymes Drosha and dicer in epithelial skin cancer. Cancer Invest. 2010;28: 649–653. doi:10.3109/07357901003630918

83. Mularoni L, Sabarinathan R, Deu-Pons J, Gonzalez-Perez A, López-Bigas N. OncodriveFML: a general framework to identify coding and non-coding regions with cancer driver mutations. Genome Biol. 2016;17: 128. doi:10.1186/s13059-016-0994-0

84. Malhotra N, Leyva-Castillo JM, Jadhav U, Barreiro O, Kam C, O’Neill NK, et al. RORα-expressing T regulatory cells restrain allergic skin inflammation. Sci Immunol. 2018;3. doi:10.1126/sciimmunol.aao6923

85. Houle AA, Gibling H, Lamaze FC, Edgington HA, Soave D, Fave M-J, et al. Aberrant PRDM9 expression impacts the pan-cancer genomic landscape. Genome Res. 2018;28: 1611–1620. doi:10.1101/gr.231696.117

86. Nicolai S, Mahen R, Raschellà G, Marini A, Pieraccioli M, Malewicz M, et al. ZNF281 is recruited on DNA breaks to facilitate DNA repair by non-homologous end joining. Oncogene. 2020;39: 754–766. doi:10.1038/s41388-019-1028-7

87. Pasparakis M, Courtois G, Hafner M, Schmidt-Supprian M, Nenci A, Toksoy A, et al. TNF-mediated inflammatory skin disease in mice with epidermis-specific deletion of IKK2. Nature. 2002;417: 861–866. doi:10.1038/nature00820

88. Stratis A, Pasparakis M, Markur D, Knaup R, Pofahl R, Metzger D, et al. Localized inflammatory skin disease following inducible ablation of I kappa B kinase 2 in murine epidermis. J Invest Dermatol. 2006;126: 614–620. doi:10.1038/sj.jid.5700092

89. Cornish GH, Tung SL, Marshall D, Ley S, Seddon BP. Tissue Specific Deletion of Inhibitor of Kappa B Kinase 2 with OX40-Cre Reveals the Unanticipated Expression from the OX40 Locus in Skin Epidermis. PLoS ONE. 2012;7. doi:10.1371/journal.pone.0032193

90. Kirkley KS, Walton KD, Duncan C, Tjalkens RB. Spontaneous Development of Cutaneous Squamous Cell Carcinoma in Mice with Cell-specific Deletion of Inhibitor of κB Kinase 2. Comp Med. 2017;67: 407–415.

91. Boyle GM, Woods SL, Bonazzi VF, Stark MS, Hacker E, Aoude LG, et al. Melanoma cell invasiveness is regulated by miR-211 suppression of the BRN2 transcription factor. Pigment Cell Melanoma Res. 2011;24: 525–537. doi:https://doi.org/10.1111/j.1755-148X.2011.00849.x

92. Simmons JL, Pierce CJ, Al-Ejeh F, Boyle GM. MITF and BRN2 contribute to metastatic growth after dissemination of melanoma. Sci Rep. 2017;7: 10909. doi:10.1038/s41598-017-11366-y

93. Zhao G, Wei Z, Guo Y. MicroRNA-107 is a novel tumor suppressor targeting POU3F2 in melanoma. Biol Res. 2020;53. doi:10.1186/s40659-020-00278-3

94. Mermel CH, Schumacher SE, Hill B, Meyerson ML, Beroukhim R, Getz G. GISTIC2.0 facilitates sensitive and confident localization of the targets of focal somatic copy-number alteration in human cancers. Genome Biol. 2011;12: R41. doi:10.1186/gb-2011-12-4-r41

95. Ikeda S, Goodman AM, Cohen PR, Jensen TJ, Ellison CK, Frampton G, et al. Metastatic basal cell carcinoma with amplification of PD-L1: exceptional response to anti-PD1 therapy. Npj Genomic Med. 2016;1: 1–5. doi:10.1038/npjgenmed.2016.37

96. Chandramouli A, Shi J, Feng Y, Holubec H, M.Shanas R, Bhattacharyya AK, et al. Haploinsufficiency of the cdc2l gene contributes to skin cancer development in mice. Carcinogenesis. 2007;28: 2028–2035. doi:10.1093/carcin/bgm066

97. de Cid R, Riveira-Munoz E, Zeeuwen PLJM, Robarge J, Liao W, Dannhauser EN, et al. Deletion of the late cornified envelope LCE3B and LCE3C genes as a susceptibility factor for psoriasis. Nat Genet. 2009;41: 211–215. doi:10.1038/ng.313

98. Myskowski PL, Pollack MS, Schorr E, Dupont B, Safai B. Human leukocyte antigen associations in basal cell carcinoma. J Am Acad Dermatol. 1985;12: 997–1000. doi:10.1016/s0190-9622(85)70127-2

99. Hua LA, Kagen CN, Carpenter RJ, Goltz RW. HLA and beta 2-microglobulin expression in basal and squamous cell carcinomas of the skin. Int J Dermatol. 1985;24: 660–663. doi:10.1111/j.1365-4362.1985.tb05719.x

100. Markey AC, Churchill LJ, MacDonald DM. Altered expression of major histocompatibility complex (MHC) antigens by epidermal tumours. J Cutan Pathol. 1990;17: 65–71. doi:10.1111/j.1600-0560.1990.tb00058.x

101. Streilein JW. Immunogenetic Factors in Skin Cancer. N Engl J Med. 1991;325: 884–887. doi:10.1056/NEJM199109193251210

102. García-Plata D, Mozos E, Carrasco L, Solana R. HLA molecule expression in cutaneous squamous cell carcinomas: an immunopathological study and clinical-immunohistopathological correlations. Histol Histopathol. 1993;8: 219–226.

103. Johnson DB, Estrada MV, Salgado R, Sanchez V, Doxie DB, Opalenik SR, et al. Melanoma-specific MHC-II expression represents a tumour-autonomous phenotype and predicts response to anti-PD-1/PD-L1 therapy. Nat Commun. 2016;7: 10582. doi:10.1038/ncomms10582

104. Chen Y-Y, Chang W-A, Lin E-S, Chen Y-J, Kuo P-L. Expressions of HLA Class II Genes in Cutaneous Melanoma Were Associated with Clinical Outcome: Bioinformatics Approaches and Systematic Analysis of Public Microarray and RNA-Seq Datasets. Diagnostics. 2019;9. doi:10.3390/diagnostics9020059

105. Stanisz H, Saul S, Müller CSL, Kappl R, Niemeyer BA, Vogt T, et al. Inverse regulation of melanoma growth and migration by Orai1/STIM2-dependent calcium entry. Pigment Cell Melanoma Res. 2014;27: 442–453. doi:https://doi.org/10.1111/pcmr.12222

106. Isvoranu G, Surcel M, Huică R-I, Munteanu AN, Pîrvu IR, Ciotaru D, et al. Natural killer cell monitoring in cutaneous melanoma - new dynamic biomarker. Oncol Lett. 2019;17: 4197–4206. doi:10.3892/ol.2019.10069

107. Sanchez-Canteli M, Hermida-Prado F, Sordo-Bahamonde C, Montoro-Jiménez I, Pozo-Agundo E, Allonca E, et al. Lectin-Like Transcript 1 (LLT1) Checkpoint: A Novel Independent Prognostic Factor in HPV-Negative Oropharyngeal Squamous Cell Carcinoma. Biomedicines. 2020;8. doi:10.3390/biomedicines8120535

108. Lone MA, Hülsmeier AJ, Saied EM, Karsai G, Arenz C, Eckardstein A von, et al. Subunit composition of the mammalian serine-palmitoyltransferase defines the spectrum of straight and methyl-branched long-chain bases. Proc Natl Acad Sci. 2020;117: 15591–15598. doi:10.1073/pnas.2002391117

109. Thomas NE, Edmiston SN, Tsai YS, Parker JS, Googe PB, Busam KJ, et al. Utility of TERT Promoter Mutations for Cutaneous Primary Melanoma Diagnosis. Am J Dermatopathol. 2019;41: 264–272. doi:10.1097/DAD.0000000000001259

110. Shaughnessy M, Njauw C-N, Artomov M, Tsao H. Classifying Melanoma by TERT Promoter Mutational Status. J Invest Dermatol. 2020;140: 390–394.e1. doi:10.1016/j.jid.2019.06.149

111. Yang E, Zha J, Jockel J, Boise LH, Thompson CB, Korsmeyer SJ. Bad, a heterodimeric partner for Bcl-xL and Bcl-2, displaces bax and promotes cell death. Cell. 1995;80: 285–291. doi:10.1016/0092-8674(95)90411-5

112. Datta SR, Dudek H, Tao X, Masters S, Fu H, Gotoh Y, et al. Akt phosphorylation of BAD couples survival signals to the cell-intrinsic death machinery. Cell. 1997;91: 231–241. doi:10.1016/s0092-8674(00)80405-5

113. Sastry KSR, Al-Muftah MA, Li P, Al-Kowari MK, Wang E, Ismail Chouchane A, et al. Targeting proapoptotic protein BAD inhibits survival and self-renewal of cancer stem cells. Cell Death Differ. 2014;21: 1936–1949. doi:10.1038/cdd.2014.140

114. Sastry KS, Ibrahim WN, Chouchane AI. Multiple signaling pathways converge on proapoptotic protein BAD to promote survival of melanocytes. FASEB J. 2020;34: 14602–14614. doi:https://doi.org/10.1096/fj.202001260RR

115. Vander Heiden MG, Li XX, Gottleib E, Hill RB, Thompson CB, Colombini M. Bcl-xL promotes the open configuration of the voltage-dependent anion channel and metabolite passage through the outer mitochondrial membrane. J Biol Chem. 2001;276: 19414–19419. doi:10.1074/jbc.M101590200

116. Danial NN, Gramm CF, Scorrano L, Zhang C-Y, Krauss S, Ranger AM, et al. BAD and glucokinase reside in a mitochondrial complex that integrates glycolysis and apoptosis. Nature. 2003;424: 952–956. doi:10.1038/nature01825

117. Seo SY, Chen Y, Ivanovska I, Ranger AM, Hong SJ, Dawson VL, et al. BAD Is a Pro-survival Factor Prior to Activation of Its Pro-apoptotic Function*. J Biol Chem. 2004;279: 42240–42249. doi:10.1074/jbc.M406775200

118. Danial NN, Walensky LD, Zhang C-Y, Choi CS, Fisher JK, Molina AJA, et al. Dual role of proapoptotic BAD in insulin secretion and beta cell survival. Nat Med. 2008;14: 144–153. doi:10.1038/nm1717

119. Roy SS, Madesh M, Davies E, Antonsson B, Danial N, Hajnóczky G. Bad targets the permeability transition pore independent of Bax or Bak to switch between Ca2+- dependent cell survival and death. Mol Cell. 2009;33: 377–388. doi:10.1016/j.molcel.2009.01.018

120. Berman SB, Chen Y, Qi B, McCaffery JM, Rucker EB, Goebbels S, et al. Bcl-x L increases mitochondrial fission, fusion, and biomass in neurons. J Cell Biol. 2009;184: 707–719. doi:10.1083/jcb.200809060

121. Aouacheria A, Baghdiguian S, Lamb HM, Huska JD, Pineda FJ, Hardwick JM. Connecting mitochondrial dynamics and life-or-death events via Bcl-2 family proteins. Neurochem Int. 2017;109: 141–161. doi:10.1016/j.neuint.2017.04.009

122. Tomková H, Fujimoto W, Arata J. Expression of the bcl-2 homologue bax in normal human skin, psoriasis vulgaris and non-melanoma skin cancers. Eur J Dermatol EJD. 1998;8: 256–260.

123. Karst AM, Dai DL, Martinka M, Li G. PUMA expression is significantly reduced in human cutaneous melanomas. Oncogene. 2005;24: 1111–1116. doi:10.1038/sj.onc.1208374

124. Aras S, Bai M, Lee I, Springett R, Hüttemann M, Grossman LI. MNRR1 (formerly CHCHD2) is a bi-organellar regulator of mitochondrial metabolism. Mitochondrion. 2015;20: 43–51. doi:10.1016/j.mito.2014.10.003

125. Liu Y, Clegg HV, Leslie PL, Di J, Tollini LA, He Y, et al. CHCHD2 inhibits apoptosis by interacting with Bcl-x L to regulate Bax activation. Cell Death Differ. 2015;22: 1035–1046. doi:10.1038/cdd.2014.194

126. Aras S. Mitochondrial autoimmunity and MNRR1 in breast carcinogenesis. 2019; 12.

127. Rosmarin AG, Resendes KK, Yang Z, McMillan JN, Fleming SL. GA-binding protein transcription factor: a review of GABP as an integrator of intracellular signaling and protein-protein interactions. Blood Cells Mol Dis. 2004;32: 143–154. doi:10.1016/j.bcmd.2003.09.005

128. Funayama M, Ohe K, Amo T, Furuya N, Yamaguchi J, Saiki S, et al. CHCHD2 mutations in autosomal dominant late-onset Parkinson’s disease: a genome-wide linkage and sequencing study. Lancet Neurol. 2015;14: 274–282. doi:10.1016/S1474-4422(14)70266-2

129. White RM, Cech J, Ratanasirintrawoot S, Lin CY, Rahl PB, Burke CJ, et al. DHODH modulates transcriptional elongation in the neural crest and melanoma. Nature. 2011;471: 518–522. doi:10.1038/nature09882

130. Garcia-Bermudez J, Baudrier L, La K, Zhu XG, Fidelin J, Sviderskiy VO, et al. Aspartate is a limiting metabolite for cancer cell proliferation under hypoxia and in tumors. Nat Cell Biol. 2018;20: 775–781. doi:10.1038/s41556-018-0118-z

131. Qian Y, Liang X, Kong P, Cheng Y, Cui H, Yan T, et al. Elevated DHODH expression promotes cell proliferation via stabilizing β-catenin in esophageal squamous cell carcinoma. Cell Death Dis. 2020;11: 1–13. doi:10.1038/s41419-020-03044-1

132. Qin J-J, Nag S, Wang W, Zhou J, Zhang W-D, Wang H, et al. NFAT as cancer target: mission possible? Biochim Biophys Acta. 2014;1846: 297–311. doi:10.1016/j.bbcan.2014.07.009

133. Mancini M, Toker A. NFAT proteins: emerging roles in cancer progression. Nat Rev Cancer. 2009;9: 810–820. doi:10.1038/nrc2735

134. Zhang X, Zhang Z, Cheng J, Li M, Wang W, Xu W, et al. Transcription Factor NFAT1 Activates the mdm2 Oncogene Independent of p53. J Biol Chem. 2012;287: 30468–30476. doi:10.1074/jbc.M112.373738

135. Griewank KG, Murali R, Schilling B, Schimming T, Möller I, Moll I, et al. TERT promoter mutations are frequent in cutaneous basal cell carcinoma and squamous cell carcinoma. PloS One. 2013;8: e80354. doi:10.1371/journal.pone.0080354

136. Scott GA, Laughlin TS, Rothberg PG. Mutations of the TERT promoter are common in basal cell carcinoma and squamous cell carcinoma. Mod Pathol Off J U S Can Acad Pathol Inc. 2014;27: 516–523. doi:10.1038/modpathol.2013.167

137. Chang ALS, Tran DC, Cannon JGD, Li S, Jeng M, Patel R, et al. Pembrolizumab for advanced basal cell carcinoma: An investigator-initiated, proof-of-concept study. J Am Acad Dermatol. 2019;80: 564–566. doi:10.1016/j.jaad.2018.08.017

138. Jiang H, Lei R, Ding S-W, Zhu S. Skewer: a fast and accurate adapter trimmer for next-generation sequencing paired-end reads. BMC Bioinformatics. 2014;15: 182. doi:10.1186/1471-2105-15-182

139. Díaz-Gay M, Vila-Casadesús M, Franch-Expósito S, Hernández-Illán E, Lozano JJ, Castellví-Bel S. Mutational Signatures in Cancer (MuSiCa): a web application to implement mutational signatures analysis in cancer samples. BMC Bioinformatics. 2018;19: 224. doi:10.1186/s12859-018-2234-y

140. Alexandrov LB, Nik-Zainal S, Wedge DC, Aparicio SAJR, Behjati S, Biankin AV, et al. Signatures of mutational processes in human cancer. Nature. 2013;500: 415–421. doi:10.1038/nature12477

141. Lawrence MS, Stojanov P, Polak P, Kryukov GV, Cibulskis K, Sivachenko A, et al. Mutational heterogeneity in cancer and the search for new cancer-associated genes. Nature. 2013;499: 214–218. doi:10.1038/nature12213

142. Cingolani P, Platts A, Wang LL, Coon M, Nguyen T, Wang L, et al. A program for annotating and predicting the effects of single nucleotide polymorphisms, SnpEff. Fly (Austin). 2012;6: 80–92. doi:10.4161/fly.19695

143. Kozlowski P, Roberts P, Dabora S, Franz D, Bissler J, Northrup H, et al. Identification of 54 large deletions/duplications in TSC1 and TSC2 using MLPA, and genotype-phenotype correlations. Hum Genet. 2007;121: 389–400. doi:10.1007/s00439-006-0308-9

144. Marcinkowska M, Wong K-K, Kwiatkowski DJ, Kozlowski P. Design and Generation of MLPA Probe Sets for Combined Copy Number and Small-Mutation Analysis of Human Genes: EGFR as an Example. Sci World J. 2010;10: 2003–2018. doi:10.1100/tsw.2010.195

145. Galka-Marciniak P, Urbanek-Trzeciak MO, Nawrocka PM, Kozlowski P. A pan-cancer atlas of somatic mutations in miRNA biogenesis genes. Nucleic Acids Res. 2021;49: 601–620. doi:10.1093/nar/gkaa1223

146. Zhou X, Edmonson MN, Wilkinson MR, Patel A, Wu G, Liu Y, et al. Exploring genomic alteration in pediatric cancer using ProteinPaint. Nat Genet. 2016;48: 4–6. doi:10.1038/ng.3466

147. UniProt Consortium. UniProt: a worldwide hub of protein knowledge. Nucleic Acids Res. 2019;47: D506–D515. doi:10.1093/nar/gky1049

148. Crowdis J, He MX, Reardon B, Van Allen EM. CoMut: visualizing integrated molecular information with comutation plots. Bioinformatics. 2020;36: 4348–4349. doi:10.1093/bioinformatics/btaa554

149. Lewis BP, Burge CB, Bartel DP. Conserved seed pairing, often flanked by adenosines, indicates that thousands of human genes are microRNA targets. Cell. 2005;120: 15–20. doi:10.1016/j.cell.2004.12.035

150. Zuker M. Mfold web server for nucleic acid folding and hybridization prediction. Nucleic Acids Res. 2003;31: 3406–3415.

151. Huang H-Y, Chien C-H, Jen K-H, Huang H-D. RegRNA: an integrated web server for identifying regulatory RNA motifs and elements. Nucleic Acids Res. 2006;34: W429–434. doi:10.1093/nar/gkl333

